# Reconstruction and variability of tropical pollination networks in the Brazilian Atlantic Forest

**DOI:** 10.1101/2021.11.22.469524

**Authors:** Juliana Pereira, Federico Battiston, Milton Cezar Ribeiro, Ferenc Jordán

## Abstract

1. Loss of biodiversity comprehends not only the extinction of individual species, but also the loss of the ecological interactions among them. Survival of species, continuation of ecosystem functioning in nature, and ecosystem services to humans depend on the maintenance of well-functioning networks of species interactions (e.g. plant-pollinator networks and food webs). Analyses of ecological networks often rely on biased and incomplete survey data, especially in species-rich areas, such as the tropics.
2. We used a network inference method to reconstruct pollination data compiled from a large tropical rainforest habitat extent. To gain insight into the characteristics of plant-pollinator interactions across the region, we combined the reconstructed pollination network with species distribution modeling to obtain local pollination networks throughout the area. We explored how global network properties relate to natural forest cover and land cover heterogeneity.
3. We found that some network properties (the sum and evenness of link weights, connectance and nestedness) are positively correlated with forest cover, indicating that networks in sites with more natural habitat have greater diversity of interactions, stability and resilience. Modularity was not related to forest cover, but seemed to reflect habitat heterogeneity, due to the broad spatial scale of the study.
4. We believe that the methodology suggested here can facilitate the use of incomplete network data in a reliable way, and allow us to better understand and protect networks of species interactions in high biodiversity regions of the world.

## Introduction

The fast decline in biodiversity currently underway has been called the sixth mass extinction (Barnosky et al., 2011). The loss encompasses not only individual species, but also ecological interactions, often at a higher rate (Valiente-Banuet et al. 2015, Kovács-Hostyánszki et al. 2019). Since many key functional aspects and services important for the maintenance of natural biomes and human subsistence depend on biotic interactions, species interactions should be treated as a major component determining the health of ecosystems (Okuyama and Holland 2008, Tylianakis et al. 2010, Valiente-Banuet et al. 2015). For example, plant-animal mutualistic networks, such as pollination and seed dispersal, sustain terrestrial biodiversity and human food security (Valdovinos, 2019). However, pollinating insects are declining in many parts of the world because of human disturbances (Machado et al., 2020), especially habitat loss (Vanbergen & the Insect Pollinators Initiative, 2013), but also landscape simplification (Dainese et al., 2019), population subdivision, and consequent changes in behavior and in interspecific interactions (Fischer & Lindenmayer, 2007). The value of ecosystem services provided globally by pollinating insects is estimated to be around 7% of the total of agricultural food production (IPBES, 2016). Around 70% of the 124 main crops consumed by humans depend on animal pollinators (Klein et al. 2007, Gallai et al. 2009), and so do most flowering plant species in the wild (NRC, 2007), especially in the tropics (Ghazoul 2005, Tokumoto 2015).

Because no species is an island, effective conservation action must protect not only individual species, but the functioning of the whole ecological community, easily represented by the web of interspecies interactions (Tylianakis et al., 2010). This focus on relationships or interactions is at the heart of Ecology as a science, and it is therefore not surprising that network tools and concepts have a long tradition in ecological studies. In fact, network ecology is a large and rapidly growing subfield within the discipline (Borrett et al., 2014). Studies dealing with species interaction networks often rely on biological survey data, sometimes originally obtained with different goals in mind. These data typically suffer from observation errors, incomplete and biased sampling, particularly when dealing with large geographic extents and species-rich systems. Network inference techniques developed in the domain of Network Science can help in the reconstruction of incomplete data (e.g. Peixoto 2018, Young et al. 2019), allowing us to capture patterns at a large scale, gain further insight into ecosystem functioning, and hopefully build better conservation strategies.

In this study, we sought to: (1) construct local plant-pollinator networks across a highly impacted tropical rainforest habitat; (2) examine some structural network properties; and (3) explore how network structure relates to natural forest cover and land cover heterogeneity. For this, we used a database on plant-insect flower visitor interactions from the Atlantic Series data papers collection. We hypothesize that the number of species, number of links, sum and evenness of link weights, connectance and modularity will increase with forest cover, while nestedness and centralization will decrease.

## Material and Methods

### Study area

The neotropical Atlantic Forest (AF) is a hotspot of biodiversity that originally covered more than 1.5 million km^2^ along the Atlantic coast of South America, including a wide variety of environmental conditions (Ribeiro et al., 2011). As a result of centuries of clearing for timber, agriculture and urban settlement, and continued anthropogenic pressure, it has been reduced to around 12% of its original extent, and subsists today almost exclusively in small fragments (< 100 ha) (Ribeiro et al., 2009). Most of the biome (64.8%) is now occupied by pasture and crops (Mapbiomas 2018 data; see Souza et al. 2020). Land cover heterogeneity is highest at areas of intermediate forest cover. The AF is a particularly good system for studying forest loss and its consequences, because it encompasses a broad gradient of fragmentation due to historical land conversion (Marjakangas et al., 2020). We divided the AF domain (sensu Ribeiro et al. 2009) within the national continental borders of Brazil in local sites (Marjakangas et al., 2020), using hexagonal grids in three scales: 25, 50 and 100 km of site length in the N-S direction, using Albers projection to ensure equal area to all sites, with SIRGAS 2000 datum, in QGIS version 3.4 (QGIS.org, 2020). To each local site (at each scale) corresponds a local plant-pollinator interaction network. We used different scales to ensure robustness of the results, since the environmental drivers that affect geographic variation in species richness can occur at, and interact across, different scales (Wilson et al. 2020, Wright et al. 2020).

### Building plant-pollinator networks

The local interaction networks corresponding to each site combine information on species interactions in general and spatial co-occurrence locally. Following the approach by Marjakangas and colleagues (2020), we first need to obtain these two types of networks separately, and then multiply them to obtain the local networks (Fig. 1). We describe these methodological steps below.

**Figure 1.**
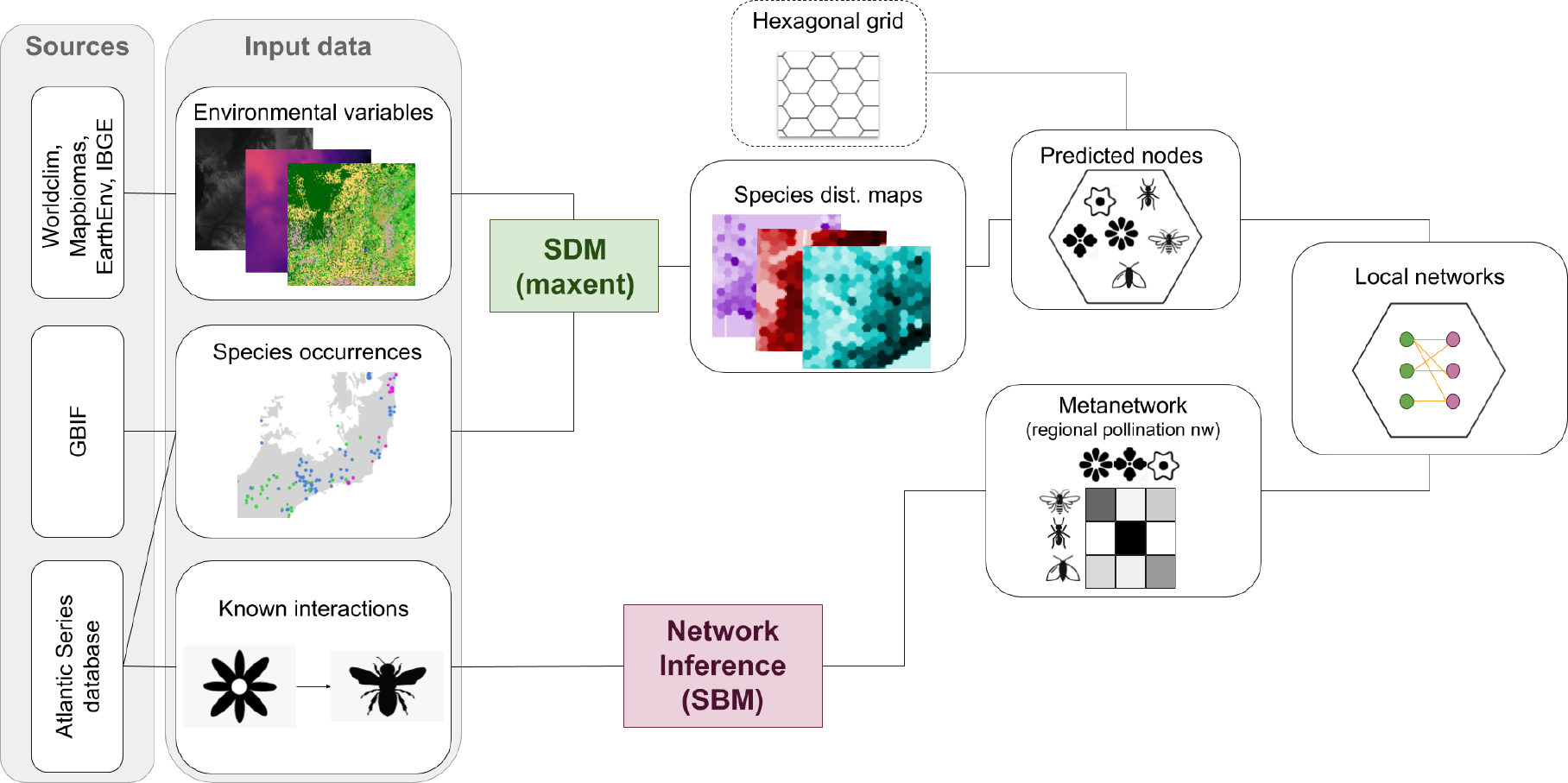
Reconstruction of local pollination networks from occurrence records of species and interactions. The study region is divided into local sites with a grid. We predict species occurrence and co-occurrence in each site using species distribution modeling, and model a regional pollination network using network inference. Multiplying co-occurrence and interaction probability networks, we obtain local pollination networks.

### Interaction network reconstruction

For the regional network of interactions, we used the Atlantic Plant-Insect database, a compilation of georeferenced interaction events observed between plants and insect flower visitors across the Brazilian AF (in prep.; see the complete Atlantic Data Papers series at https://esajournals.onlinelibrary.wiley.com/doi/toc/10.1002/(ISSN)1939-9170.AtlanticPapers and at https://github.com/LEEClab/Atlantic_series). We kept only observations with classification at the species level, within the Brazilian territory and the AF domain. This comprised our initial regional pollination network, or *metanetwork*, with 1273 species (nodes) and 2895 unique interaction pairs (links).

As is the case with most empirical networks, sampling of interactions in biological surveys is incomplete in our dataset. Studies show that sampling bias tends to strongly underestimate the number of interactions and overestimate the degree of specialization (Trøjelsgaard & Olesen, 2013; Young et al., 2019). To obtain a more reliable picture of pollination interactions across the AF, we applied a method of network reconstruction based on stochastic blockmodels to our metanetwork. Blockmodels are generative models for networks, where nodes are classified into blocks according to their profile of interactions (i.e. nodes that connect or not to the same groups of other nodes are put in the same group). Probabilities are used to describe interactions between blocks. For example, let’s imagine a situation where white flowers usually are visited by moths, and never by butterflies, pink flowers are visited by both butterflies and bees, and yellow and blue flowers are visited only by bees. The blockmodel corresponding to this network would have six blocks: moths, butterflies, bees, yellow and blue flowers, white flowers, pink flowers. We visualize the model as a block matrix (Supplementary material Fig. 1), with entry values *b_ij_* corresponding to the probability of an interaction between a node of block *i* and a node of block *j*.

A block matrix like this can be used to draw network samples. Conversely, we can fit a block matrix to a known incomplete network and, as a second step, infer missing links and network structure based on the blocks and probabilities, in order to reconstruct the network. We did this using the python package graph-tool (Peixoto, 2014), which uses Bayesian inference to extract the underlying optimal block structure. First, we fitted a nested stochastic blockmodel to our binary metanetwork. Next, we reconstructed our metanetwork using the fitted blockmodel as prior. We used the function UncertainBlockState, adequate for cases in which we do not have repeated measurements of each link, but can offer extraneous error estimates as independent edge probabilities (Peixoto, 2014). This applies well to our case, because the data is a compilation from many different studies, with different geographical extents, methods of survey and research questions, so that the numbers of observations recorded for each pair are not comparable. We used three error estimates: first, for all forbidden links (i.e. plant-plant or pollinator-pollinator links), we set the uncertainty value to 0, indicating our previous knowledge that these links don’t exist. Secondly, we must choose an uncertainty value for the observed links. After testing different values ranging between 0.75 and 0.99, we chose the value 0.98, because it resulted in the best reconstructed network (no forbidden links were created, and the maximum number of the original links was kept). Thirdly, for each non-edge (non observed plant-pollinator links), we picked a value chosen to preserve the expected density of the original network (see Supplementary Material for the code and further details). The resulting reconstructed network had 49406 links, none of which were forbidden, and had lost only 2 of the original edges. Each resulting link has a weight corresponding to its posterior probability (e.g. the probability that this edge exists), which we interpret as the potential of interaction between species.

### Co-occurrences and local networks

Next, we performed species distribution modeling (Guisan and Zimmermann 2000) with Maxent (Phillips et al., 2004) to predict the occurrences of species in each site, using the R package ENMeval (Muscarella et al., 2014). As labeled data, we included the georeferenced interaction points from the pollination dataset, complemented by additional species occurrence points from the Global Biodiversity Information Facility (dataset available at DOI10.15468/dl.by6nuj: GBIF.org 2018) filtered with the R package CoordinateCleaner (Zizka et al., 2019). We removed any point records of the same species located within the same 1 km x 1 km raster cell of environmental covariates used in the model to avoid causing bias in the distribution modeling. Only species with at least 5 instances were included in the analysis. The environmental factors reported to affect plant and insect distributions the most are temperature, precipitation and soil type (Savopoulou-Soultani et al. 2012, Khaliq et al. 2014, Ding et al. 2016, Ali et al. 2019). For this reason, we used as explanatory environmental variables topographic, soil, climatic and land cover covariates, with final resolution of 1 km (Supplementary Material Table 1). For background points, we used the weighted-target group approach (i.e. occurrence points of all plant or pollinator species other than the one being modeled), which reduces the influence of sampling bias (Anderson 2003, Muscarella et al. 2014). We selected models with ΔAICc = 0, and secondarily with maximum AUC on testing data. As final output, we chose the cumulative output maps, which can be interpreted in terms of omission rate, i.e. thresholding the map at a value *c* to binarize it into prediction of presence or absence of the species would omit approximately *c*% of presences (Merow et al., 2013). We discarded models that performed poorly (AUC < 0.7), and were left with 389 species (297 plants and 92 pollinators, Supplementary Material Table 2). Of these, 223 species’ distributions are positively correlated with natural forest cover (%). Since the most generalist species (*Apis mellifera*, *Trigona spinipes*, *Bombus morio* and *Plebeia droryana*) did not model well, we had to exclude them. Their absence in our networks could considerably alter their structure and mask some of the properties’ correlations we expected to find. Therefore, we decided to proceed with the analysis considering, in parallel, networks with all 389 species (all-spp networks) and networks only with the 223 forest species (forest-spp networks), and compare the results at the end, considering that, broadly speaking, it is species and networks associated with forests we would be most interested in protecting in the AF.

From the SDM maps of cumulative occurrence probability, we computed the average occurrence probability of each species in each site, at each scale. In order to interact, two potentially interacting species must co-occur in space (Tylianakis et al., 2010). Therefore, we multiplied the average occurrence probability of all plant-pollinator pairs for each site, to obtain the local probabilities of co-occurrence. Next, we multiplied the probability of co-occurrence of each plant-pollinator pair in each site by the probability of interaction of the pair according to our pollination metanetwork. In this way, we obtained local pollination networks for each site, with probabilistic link weights, at each scale. We did this including all the 389 species for which SDM was successful (all-spp networks), and then repeated it including only the species whose spatial distribution is positively correlated (Spearman’s ρ >0 with *P* < 0.05) with natural forest cover (forest-spp networks).

### Linking network properties with forest and land cover heterogeneity

For each scale, for all-spp and forest-spp networks, we computed measures that provide a quantification of network properties relevant to particular ecological functions: (1) number of plant and pollinator nodes, or species (N_pl_, N_po_); (2) number of links (L); (3) sum of link weights, informing about the general intensity of interactions in the community (SLW); (4) evenness of node strength for plants and pollinators, measured by Shannon entropy (HS_pl_, HS_po_), where node strength is the sum of the weights of the links connected to it; (5) weighted connectance or density, which quantifies how many links are present, in comparison to all possible links, taking weights into account (C); (6) centralization, which shows how much the links are centralized on one or a few nodes, based on eigenvector centrality (Centr; Freeman 1978); (7) bipartite modularity, which shows how much the network is compartmented into dense groups of nodes called modules, using the method by Beckett (2016) (M); (8) number of modules (Nb_mod_); (9) assortativity based on node strength, which shows how much nodes tend to connect with others of similar strength (A); and (10) nestedness, which shows how much the neighborhoods of specialist species are subsets of the neighborhoods of generalists, using the spectral radius index and row column totals average null model (Nest; Beckett et al. 2014, Mariani et al., 2019). The use of this null model for the computation of nestedness is important to capture only nestedness beyond what is explained by degree (Lewinsohn et al., 2006).

Finally, we tested the correlation of each of these network properties with forest cover (%) and land cover heterogeneity (HLC, Shannon entropy of land cover categories per site) using Spearman’s rank correlation. These computations were made using the packages igraph (Csardi & Nepusz, 2006), bipartite (Dormann et al., 2008), vegan (Oksanen et al., 2019) and Falcon (Beckett et al., 2014) in R (R Core Team, 2020) with RStudio (RStudio Team, 2019).

## Results

Investigating our local pollination networks at the most detailed scale (25 km), we found that degree and node strength were positively correlated (mean Spearman’s ρ = 0.57 and 0.56 for all-spp and forest-spp networks, respectively) in agreement with Bascompte et al. (2006). Most link weights were small (average link weight between 0.01 and 0.22 for all-spp and between 0.02 and 0.36 for forest-spp networks). Connectance and assortativity were always below 0.04 (Supplementary Material Table 3), while centralization was between 0.75 and 0.80 for all-spp and 0.66-0.74 for forest-spp networks. Modularity was always positive, and the majority of modules are nested (75.07% in all-spp and 75.04% in forest-spp networks, on average). Nestedness was not correlated correlated with the number of links (*P* = 0.18 and Spearman’s ρ = −0.07 for all-spp and forest-spp, respectively).

We observed similar correlations between network properties and forest cover or land cover heterogeneity, independently of scale (Supplementary Material Table 4 and Figure 2). In some cases, correlations became stronger with increasing site area (e.g. SLW ~ forest cover in forest-spp networks), while in other cases they became weaker (e.g. connectance ~ forest cover in forest-spp network). In most instances, the tendencies were also similar between all-spp and forest-spp networks. We observe clearer tendencies in the forest-spp networks: network size (N and L) and centrality appear particularly related to land cover heterogeneity, while SLW, HS, connectance, nestedness and assortativity are shown to be more related to forest cover. These patterns are not very clear when we look at all-spp networks. Modularity and the number of modules showed weak to no correlation in all cases, although modularity seems more related to land cover heterogeneity than to forest cover, when looking at forest-spp networks.

**Figure 2.**
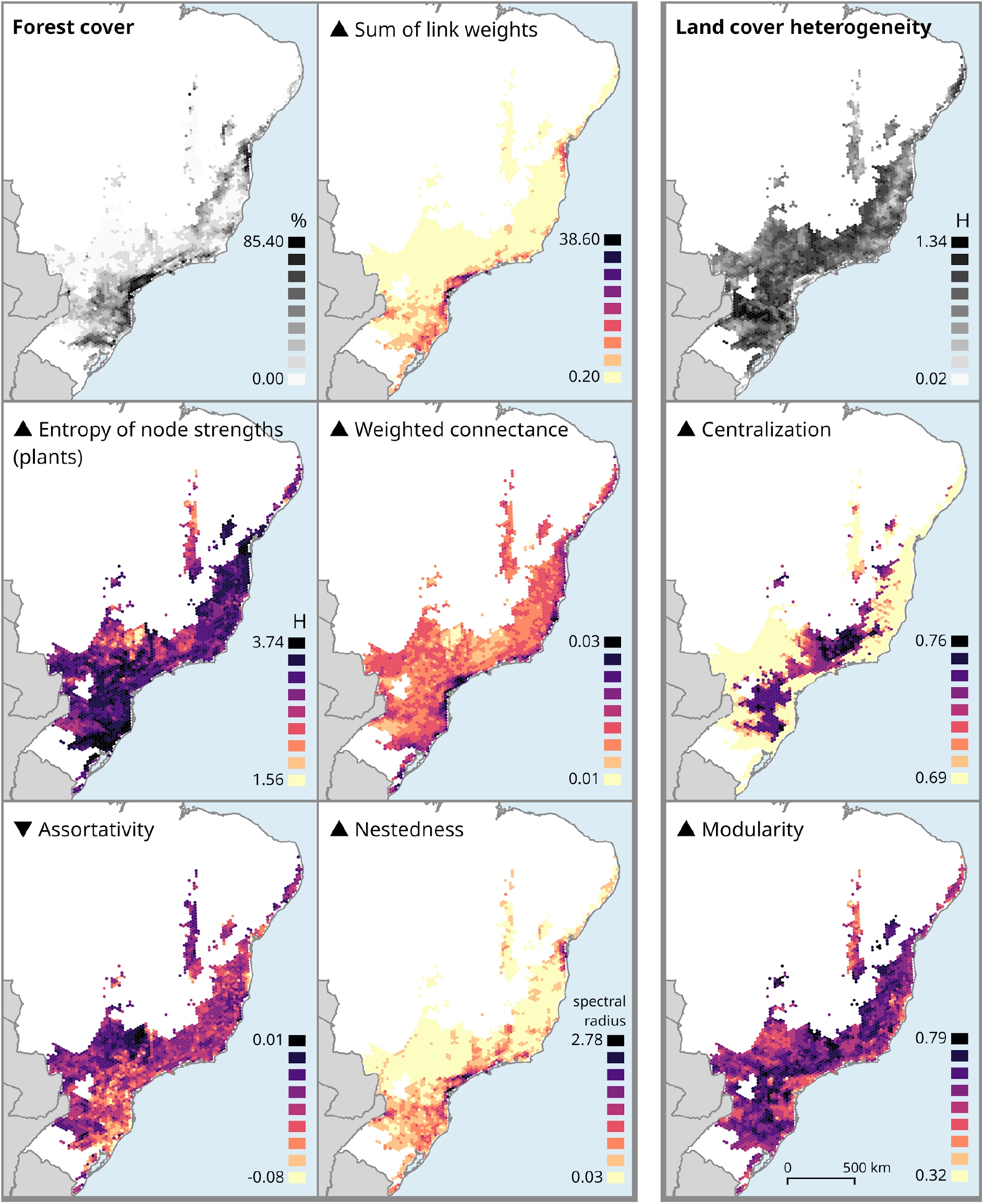
Pollination network properties correlate to habitat variables. Some network properties are more strongly correlated with forest cover (left) and others with land cover heterogeneity (right), positively (▲) or negatively (▼). Networks of forest species on sites of 25 km are shown. Results for 50 and 100 km were similar. Equal interval classes are used for the coloring.

We concentrate now on the average correlation across the different scales for each situation (Table 1). All-spp networks are smaller (N and L) both with increasing forest cover and land cover heterogeneity. Forest-spp networks’ sizes seem more related to HLC, and the smaller size is only due to fewer insect species. Actually, we observed that networks of high forest cover sites are always relatively large (N, L) and have higher S, while those in sites with less forest cover have a wide range of network sizes, but their total strength S is towards the lower end (Supplementary Material Fig. 3). In other words, only high forest cover sites are characterised by networks with higher S, while L is basically independent of forest cover. In fact, looking at forest-spp networks, it is clear that high forest cover sites have networks with stronger (although not necessarily more numerous) links (SLW and weighted connectance), as well as more evenly distributed link strengths (HS). Nestedness increases and assortativity decreases with forest cover, for forest-spp networks. No clear trend is visible for all-spp networks. Centralization, on the other hand, seems more associated with HLC (Table 1, Fig. 2).

**Table 1.** Network properties’ correlation with forest cover and land cover heterogeneity were independent of spatial scale. Forest-species-only networks exhibited a clearer pattern, with network size and centralization dependent on land cover heterogeneity, and node strength, connectance and nestedness dependent on forest cover.

| | | $N_{pl}$ | $N_{po}$ | L | SLW | $HS_{pl}$ | $HS_{po}$ | C | Centr | M | A | Nest |
| --- | --- | --- | --- | --- | --- | --- | --- | --- | --- | --- | --- | --- |
| all-spp | Forest cover | -0.35<br>▼ | -0.35<br>▼ | -0.36<br>▼ | 0.27<br>■▲ | 0.09<br>■ | 0.32<br>▲ | 0.19<br>■▲ | 0.36<br>▲ | -0.12<br>■▼ | n.s.<br>■ | 0.22<br>■▲ |
|  | HLC | -0.08<br>■ | -0.46<br>▼ | -0.44<br>▼ | -0.14<br>■▼ | n.s.<br>■ | n.s.<br>■ | n.s.<br>■ | 0.46<br>▲ | n.s.<br>■ | n.s.<br>■ | -0.09<br>■ |
| forest-spp | Forest cover | n.s.<br>■ | -0.27<br>■▼ | -0.25<br>■▼ | 0.78<br>▲ | 0.58<br>▲ | 0.56<br>▲ | 0.33<br>▲ | 0.21<br>■▲ | 0.17<br>■▲ | -0.60<br>▼ | 0.66<br>▲ |
|  | HLC | n.s.<br>■ | -0.47<br>▼ | -0.48<br>▼ | 0.18<br>■▲ | 0.16<br>■▲ | 0.12<br>■▲ | -0.17<br>■▼ | 0.47<br>▲ | 0.25<br>■▲ | -0.11<br>■▼ | 0.10<br>■ |

$N_{pl}/N_{po}$ : number of plant / pollinator species, L: number of links, SLW: sum of link weights, $HS_{pl}/HS_{po}$ : entropy of plant / pollinator node strengths, C: weighted connectance, Centr: centralization based on eigenvector centrality, M: bipartite modularity, A: assortativity, Nest: spectral radius index of nestedness. Values are average Spearman's rho correlation index across three scales (site N-S length of 25, 50 and 100 km) for all-spp and forest-spp networks. The symbols show positive i.e. $\rho > 0.3$ (▲), negative i.e. $\rho < -0.3$ (▼), weak positive i.e. $0.1 < \rho < 0.3$ (■▲), weak negative i.e. $-0.3 < \rho < -0.1$ (■▼) or no correlation i.e. $-0.1 < \rho < 0.1$ or $P > 0.05$ (■). When at least two out of the three correlations were non-significant, we show them as *n.s.* When one of the three was non-significant, we show the smallest absolute significant value. Cases of opposite correlation between all-spp and forest-spp are highlighted in grey.

There are two cases in which forest-spp and all-spp networks show opposite results, both involving only very weak correlations. First, modularity seems to decrease with forest cover for all-spp and increase for forest-spp networks. We should however keep in mind that, for forest-spp, modularity was more associated with HLC than with forest cover (Supplementary Material Table 4). Secondly, SLW in function of HLC decreases for all-spp, but increases for forest-spp. That is, in homogeneous sites, networks of forest species have weaker links, and networks of all-spp have stronger links. It is interesting to note that most homogeneous sites in the study area are occupied by agriculture rather than forest.

## Discussion

In general, sites with large and continuous natural forest host more species and interaction links than deforested or fragmented areas (Marjakangas et al., 2020). Yet, in order to better understand the functioning of both natural and human-impacted forest communities, we need to better understand the principles of their interaction networks, and whether they contain fewer species or interactions. Since our local networks are the result of multiplying two networks with probabilistic links (co-occurrence and potential of interaction), the weights of most links are extremely small, while nodes’ strengths and link weights are rarely equal to zero, making both N and L very high. Especially in sites with little forest cover, we have an enormous amount of links with weight very close to zero. Therefore, although the number of nodes and links seem to decrease with forest cover, these values carry little meaning in our context. Instead, it is by looking at node strengths (SLW and HS) that we can observe relevant patterns. Indeed, network size may be strongly affected by sampling effort (Ings et al. 2009, Trøjelsgaard and Olesen 2013), and should be regarded with care. Moreover, habitat disturbance may affect link weights more than species richness, as observed for host-parasitoid networks (Tylianakis et al., 2007). Link weights play an important role in stability, with many weak and few strong links, as we observe here, being commonly observed and leading to stable but potentially complex webs (Jordano 1987, Ings et al. 2009). We have found the sum of link weights to be higher in more forested sites. Loss of interaction links is generally expected where there is loss of habitat, although there are exceptions due to rewiring (Valiente-Banuet et al., 2015). We have also found node strengths to be more evenly distributed in sites with high forest cover, in agreement with Tokumoto (2015). This may be due to the fact that networks in degraded or fragmented areas rely more heavily on super-generalist species such as the honeybee *Apis mellifera* (Rathcke and Jules 1993, Kovács-Hostyánszki et al. 2019), which have disproportionately high node degree and strength, through establishing a large number of weaker links, and thus increasing interaction asymmetry (Tylianakis et al., 2010). The centralization index did not indicate the same tendency, being instead larger for high land cover heterogeneity. However, the exclusion of the most generalist species from the analysis may be responsible for masking network centralization in sites with less forest cover. We have nevertheless observed high absolute values of network centralization, expected in pollination webs (Martín González et al., 2010), especially in the all-spp networks, indicating that species indifferent to or negatively associated with forest increase network centralization in our system.

Networks in more forested areas had also higher density or connectance, in agreement with Morales and Vázquez (2008). High connectance is reportedly associated with stability in mutualistic networks, both in the sense of robustness to the loss of species and fast return to equilibrium after perturbations (Thebault & Fontaine, 2010). Connectance may be strongly influenced by sampling effort (Ings et al., 2009), but we could observe a clear correlation, possibly due to network reconstruction.

Nestedness is very common in plant-animal mutualistic networks (Bascompte and Jordano 2007, Trøjelsgaard and Olesen 2013), especially when coevolutionary selection is weak (Segar et al., 2020). Some studies report nestedness to be robust to sampling effort (Bascompte and Jordano 2007, Ings et al. 2009), although there are exceptions (Trøjelsgaard & Olesen, 2013). Nestedness has been reported to increase with the number of links in a study with binary networks (J. Bascompte et al., 2003). Conversely, we did not observe any correlation between these network properties, possibly because we used weighted measures. Nestedness reflects a network structure where many specialist species are reliant on a few persistent generalist species (Segar et al., 2020), and may be generated by differential dispersal, abundance, spatial distribution or similar ecological processes (Lewinsohn et al., 2006). An advantage of nestedness is that it facilitates successful responses to perturbation (Jordano, 1987): in a nested network, it is unlikely that a species ends up isolated, unless it is a very poorly connected species, whose loss will not strongly affect the functioning of the network (Tylianakis et al., 2010). Moreover, the pattern of generalist-with-specialist links provides ways for rare species to persist for a longer time (Bascompte et al. 2003, Memmott et al. 2004, Fortuna and Bascompte 2006, Thebault and Fontaine 2010). We found a clear relationship between nestedness and forest cover only when considering forest-spp only. Some studies suggest that nested networks, being more resilient, are associated with areas that suffered high impact or very variable climate (Lewinsohn et al. 2006, Tylianakis et al. 2010, Dalsgaard et al. 2013), while others report the opposite. For example, Dupont and collaborators (2009) found pollination networks in lower latitudes to be more nested. We found nestedness to increase with forest cover, agreeing with the latter. In any case, the AF biome as a whole has suffered severe anthropogenic disturbance, and even sites that still retain a large proportion of natural forest today are affected by environmental degradation and fragmentation, and host more resilient communities which are quite different from those of pristine areas (Marjakangas et al., 2020). Furthermore, in mutualistic assemblages of low specificity and high diversity we may expect compartments with nested structure (Lewinsohn et al., 2006), which is precisely what we found.

Plant-animal mutualistic networks have been reported to be asymmetric, or disassortative, which makes the entire network more resistant to secondary extinctions (Jordano 1987, Bascompte et al. 2006). One study found mutualistic networks to be more disassortative where there is more connected natural habitat (Tokumoto, 2015), and another where habitat is more fragmented (Morales & Vázquez, 2008). Our results are in line with the former. Disassortativity is also correlated with nestedness (Tylianakis et al. 2010, Jonhson et al. 2013), as we have observed.

For mutualistic networks, modularity is associated with instability, because it means reduced functional redundancy, increasing the chances of secondary extinctions (Dalsgaard et al., 2013). Nevertheless, while one-to-one specialization is extremely rare in free living species (Jordano, 1987), more specificity in large networks may lead to the formation of separate groups, leading to high modularity (Olesen et al. 2007, Ings et al. 2009, Trøjelsgaard and Olesen, 2013). Three main explanations have been proposed for modular structure in mutualistic networks. First, modular networks may, in view of this structural fragility, be associated with areas of great environmental stability, which have been left undisturbed for long periods (Dalsgaard et al., 2013). This is because, in such areas, modules corresponding to coevolutionary units with convergent traits (e.g. pollination syndromes; Jordano 1987, Olesen et al. 2007, Segar et al. 2020) and/or phylogenetically close species (Bascompte and Jordano 2007, Ings et al. 2009, Chamberlain et al. 2014) have had the opportunity to evolve unperturbed (Dalsgaard et al., 2013). Tropical habitats are an example of that, with their slow Quaternary climate change, and rich networks associated with high specialization (Dalsgaard et al., 2011; Trøjelsgaard and Olesen, 2013). In accordance with this, we found relatively high modularity values in our networks. However, we did not find a correlation between modularity and the quality (forest cover) of sites, probably because of the history of high environmental impact in the region, affecting even the most preserved habitat fragments. Secondly, modular structures may be due to the loss of native supergeneralists (Tylianakis et al., 2010), possibly caused by the presence of highly competitive alien species like the honeybee (Valido et al., 2019). Although we know the honeybee and a few native supergeneralists are present in our study area, we were unable to include these species in our analysis because of poor performance of SDM modeling, probably due to their ubiquity. Finally, modularity may also reflect habitat heterogeneity (Tokumoto, 2015), divergent selection regimes, and phylogenetic splits or clustering of closely related species (Olesen et al., 2007), giving rise to compartments that are more perceptible at a large-scale or regional level (Lewinsohn et al., 2006). Due to the coarse spatial scale of our study, it is likely that the modules we found are best accounted for by habitat heterogeneity (e.g. species associated with forest and farmland species coexisting in the same site, but in fairly separate parts of the network). Indeed, the highest correlation we obtained for modularity was with land cover heterogeneity. Concerning modularity and forest cover, we have seen contrasting results: they are negatively correlated for all-spp networks and positively for forest-spp only. This could be an indication that the non-forest species create bridges between modules of forest species in forested sites, decreasing modularity, and create modules of their own in other sites.

Nestedness and modularity may show opposite geographical patterns (Dalsgaard et al. 2013, Segar et al. 2020), but they are not necessarily mutually exclusive, appearing together especially in large (>150 species) networks (Lewinsohn et al. 2006, Ings et al. 2009, Trøjelsgaard and Olesen 2013, Mello et al. 2019). Often modular networks may be nested as a whole, and have modules that are themselves nested (Lewinsohn et al., 2006), as we have observed. Such networks may be structured as a core of generalist species that connect modules to each other (Olesen et al., 2007). This structure can be a result of specializations that confine species in modules, while within modules nestedness can emerge (Lewinsohn et al., 2006).

Our work sought to propose a methodological strategy to reconstruct interactions networks in high-diversity regions with incomplete sampling, and to present an example case study exploring the relationship between network structure, forest cover and land cover heterogeneity in the Atlantic Forest. Our results call for some considerations and suggest some avenues for future investigations. First, pollination networks are highly dynamical in species composition and interactions through time. Interactions may also change with location (Lewinsohn et al., 2006), depending on the assemblage and conditions of each habitat site. Similarly, rewiring may occur in ecological networks, increasing resilience even in the face of species loss (Vizentin-Bugoni et al., 2020). However, the structure of the network (e.g. connectance, nestedness, modularity, centralization) remains fairly constant (Dupont et al., 2009). Therefore, although in this study we have not examined how networks change over time, the results concerning network structure should still be valid. Moreover, we used probabilistic links along with network reconstruction, which should signal the possibility of connections that did not exist at the time or place of sampling, but may arise via rewiring. In further investigations involving network dynamics, node and module level properties, time series network data would be most valuable, and we would encourage field surveys in this direction. Secondly, our networks are based on potential species distribution, i.e. they are built on abiotic factors, such as climate, land cover and topographic variables. An important direction for future work is to include species interactions themselves and their phylogenetic history in the modeling of species distribution (Carnaval et al., 2014), leading to more realistic networks. Finally, flower visitation webs, as the ones used here, may give an inaccurate impression of pollination interactions, because flower visitors can exhibit specialist behavior when collecting pollen but generalist behavior when collecting nectar (Ings et al., 2009). New sampling techniques and molecular methods may hopefully bring a breakthrough here with better data.

## Conclusion

Networks of species interactions need to be targeted for conservation, because the long-term survival of species in the wild, as well as ecosystem functioning and services, depend on biotic interactions. In species-rich regions like the tropics, complete and detailed network data over large geographic extents is often rare. We have shown how network reconstruction, combined with species distribution modeling, can be used to infer local interaction networks in a tropical rainforest habitat. We have examined how some global network properties correlate with forest cover and land cover heterogeneity, finding that nestedness and the sum of interaction strengths are strongly associated with greater forest cover. We believe that our results bring further insight into the structure of pollination networks across the Atlantic Forest, which is important for conservation planning focused on ecological interactions. We hope our work can be helpful for further studies aiming to understand and protect species interaction networks, particularly in regions of the world where the very wealth of biological diversity that motivates conservation action also makes it difficult to obtain complete data.

## Acknowledgments

We are thankful to Tiago Peixoto for important assistance and advice in network reconstruction.

## Author’s contributions

All authors participated in the conception of the idea; JP and FB designed methodology; MR compiled the data; JP analysed the data; JP, FB and FJ led the writing of the manuscript. All authors contributed to the drafts and gave final approval for publication.

## Notes

### Competing Interest Statement

The authors have declared no competing interest.

### Summary of Updates

Author order modified

