## Supplementary Material for "Reconstruction and variability of tropical pollination networks in the Brazilian Atlantic Forest"

### Code

Available at https://github.com/juliana-p/Pollination_network_reconstruction.

### Table 1.

Environmental covariates used in Maxent Species Distribution Modeling of plant and pollinator species.

| **Category** | **Layers** | **Data type** | **Resolution / Scale** | **CRS (EPSG)** | **Source** | **Date of access** |
|---|---|---|---|---|---|---|
| Topography | - Elevation<br>- Eastness | Raster | 1 km | 4326 | EarthEnv | 20 Sep 2019 |
| Soil | - Soil 37 classes<br>- Derived soil 5 classes | Vector<br>Raster | 1:5.000.000<br>1 km | 4618<br>4326 | IBGE-EMBRAPA 2001<br>Reclassification of soil classes layer. | 20 Sep 2019 |
| Climatic | - BIO5: temperature seasonality (SD)<br>- BIO14: precipitation of driest month | Raster | 0.5 degrees (~ 1km) | 4326 | WorldClim with the raster R package | 20 Sep 2019 |
| Land cover | - Land cover 19 classes<br><br>- Derived density for selected classes: natural forest, planted forest, pasture, annual and perennial crop, semi-perennial crop, mosaic of agriculture and pasture, urban infrastructure<br>- Derived land cover heterogeneity | Raster<br><br>Raster<br><br>Raster | 30 m<br><br>1 km<br><br>1 km | 4326<br><br>4326<br><br>4326 | Mapbiomas 2018 with user-toolkit<br>Percentage cover<br><br>Shannon entropy index of land cover classes | 5 Dec 2019 |

**Table 2.**

Studied species, maxent SDM model performance and correlation of species distribution with natural forest (Spearman's rho). Species with AUC < 0.7 were discarded. Out of 389 successful SDM species, 223 are positively correlated with forest (forest species) and were included in the forest-spp networks.

| species | mode | AUC | rho | forest species |
|---|---|---|---|---|
| *Adenostemma brasilianum* | plant | 0.85732 | 0.69248 | yes |
| *Aechmea ornata* | plant | 0.80139 | 0.31504 | yes |
| *Albizia polycephala* | plant | 0.77779 | -0.26892 | no |
| *Alchornea sidifolia* | plant | 0.77385 | 0.52316 | yes |
| *Anadenanthera colubrina* | plant | 0.76709 | -0.44734 | no |
| *Anemopaegma scabriusculum* | plant | 0.94016 | -0.30940 | no |
| *Annona monticola* | plant | 0.84137 | -0.16885 | no |
| *Anthurium affine* | plant | 0.76552 | -0.29507 | no |
| *Aristolochia gigantea* | plant | 0.80977 | -0.27364 | no |
| *Aspilia foliosa* | plant | 0.93320 | -0.19680 | no |
| *Aspidosperma olivaceum* | plant | 0.72090 | 0.25951 | yes |
| *Aspidosperma pyrifolium* | plant | 0.84907 | -0.34950 | no |
| *Austroeupatorium inulaefolium* | plant | 0.71106 | 0.47481 | yes |
| *Baccharis anomala* | plant | 0.83243 | 0.46249 | yes |
| *Baccharis articulata* | plant | 0.87959 | 0.38370 | yes |
| *Baccharis concinna* | plant | 0.93323 | -0.10886 | no |
| *Baccharis conyzoides* | plant | 0.82988 | 0.26209 | yes |
| *Baccharis crispa* | plant | 0.80471 | 0.39800 | yes |
| *Baccharis leucopappa* | plant | 0.85564 | 0.38666 | yes |
| *Baccharis megapotamica* | plant | 0.93611 | 0.30159 | yes |
| *Baccharis* | plant | 0.84560 | 0.46777 | yes |

| | | | | |
|---|---|---|---|---|
| *microdonta* | | | | |
| *Baccharis patens* | plant | 0.82339 | 0.26512 | yes |
| *Baccharis pentodonta* | plant | 0.90453 | 0.28197 | yes |
| *Baccharis retusa* | plant | 0.80000 | 0.24422 | yes |
| *Baccharis sagittalis* | plant | 0.73929 | 0.53432 | yes |
| *Bactris setosa* | plant | 0.73994 | 0.37671 | yes |
| *Baccharis singularis* | plant | 0.74313 | 0.26816 | yes |
| *Baccharis spicata* | plant | 0.80299 | 0.27699 | yes |
| *Banara guianensis* | plant | 0.80629 | -0.09056 | no |
| *Bathysa australis* | plant | 0.75929 | 0.70472 | yes |
| *Bixa orellana* | plant | 0.71035 | -0.41092 | no |
| *Blutaparon portulacoides* | plant | 0.86194 | -0.08376 | no |
| *Buchnera longifolia* | plant | 0.84228 | n.s. | no |
| *Byrsonima correifolia* | plant | 0.92977 | -0.21843 | no |
| *Byrsonima dealbata* | plant | 0.81982 | -0.23386 | no |
| *Byrsonima gardneriana* | plant | 0.80323 | -0.37239 | no |
| *Caesalpinia echinata* | plant | 0.71704 | -0.15511 | no |
| *Caladium bicolor* | plant | 0.78934 | -0.12485 | no |
| *Calliandra brevipes* | plant | 0.70165 | 0.44358 | yes |
| *Calea harleyi* | plant | 0.87851 | -0.05176 | no |
| *Calea pinnatifida* | plant | 0.72837 | 0.43053 | yes |
| *Calliandra tweedii* | plant | 0.71713 | 0.15189 | yes |
| *Campylocentrum aromaticum* | plant | 0.76093 | 0.40335 | yes |
| *Campomanesia aurea* | plant | 0.82524 | 0.15059 | yes |
| *Camptosema coriaceum* | plant | 0.91277 | -0.15384 | no |

| | | | | |
|---|---|---|---|---|
| *Campomanesia dichotoma* | plant | 0.84135 | -0.28104 | no |
| *Campuloclinium macrocephalum* | plant | 0.78259 | 0.29776 | yes |
| *Campomanesia phaea* | plant | 0.80224 | 0.27046 | yes |
| *Camellia sinensis* | plant | 0.77557 | 0.61598 | yes |
| *Cambessedesia wurdackii* | plant | 0.90879 | -0.00276 | no |
| *Canavalia rosea* | plant | 0.88121 | -0.12914 | no |
| *Cardiospermum corindum* | plant | 0.87497 | -0.33855 | no |
| *Cassia fistula* | plant | 0.78818 | -0.09777 | no |
| *Cattleya elongata* | plant | 0.92756 | -0.07616 | no |
| *Centrolobium tomentosum* | plant | 0.77003 | -0.15676 | no |
| *Cestrum strigillatum* | plant | 0.75293 | 0.01736 | yes, but isolated in forest-spp only networks |
| *Chamaecrista calycioides* | plant | 0.74854 | 0.18861 | yes |
| *Chamaecrista desvauxii* | plant | 0.73894 | -0.25211 | no |
| *Chamaecrista rotundifolia* | plant | 0.70151 | -0.39879 | no |
| *Chamaecrista zygophylloides* | plant | 0.83682 | -0.13347 | no |
| *Chromolaena ascendens* | plant | 0.78264 | 0.50982 | yes |
| *Chromolaena congesta* | plant | 0.87109 | 0.17675 | yes |
| *Chrysolaena flexuosa* | plant | 0.89860 | 0.28989 | yes |
| *Chromolaena hirsuta* | plant | 0.88251 | 0.41729 | yes |
| *Cirrhaea dependens* | plant | 0.82905 | 0.18572 | yes |
| *Coccoloba brasiliensis* | plant | 0.88564 | -0.08002 | no |

| | | | | |
|---|---|---|---|---|
| *Coccocypselum erythrocephalum* | plant | 0.92599 | 0.00266 | yes |
| *Cocos nucifera* | plant | 0.77516 | -0.24149 | no |
| *Cochlospermum vitifolium* | plant | 0.80773 | -0.35856 | no |
| *Coffea arabica* | plant | 0.73119 | -0.03327 | no |
| *Collaea cipoensis* | plant | 0.97374 | -0.1549 | no |
| *Collaea stenophylla* | plant | 0.80686 | 0.27012 | yes |
| *Condea fastigiata* | plant | 0.72908 | 0.34177 | yes |
| *Croton blanchetianus* | plant | 0.93988 | -0.39192 | no |
| *Croton campestris* | plant | 0.89408 | -0.26980 | no |
| *Crocosmia crocosmiiflora* | plant | 0.77648 | 0.39145 | yes |
| *Croton glutinosus* | plant | 0.91573 | -0.08715 | no |
| *Croton gnaphalii* | plant | 0.97771 | 0.06815 | yes |
| *Cucurbita pepo* | plant | 0.72252 | -0.06588 | no |
| *Cuphea ericoides* | plant | 0.87612 | -0.19785 | no |
| *Cuphea flava* | plant | 0.81667 | -0.31329 | no |
| *Cuphea glutinosa* | plant | 0.91507 | 0.25684 | yes |
| *Cyclopogon apricus* | plant | 0.87219 | 0.25203 | yes |
| *Cypella amplimaculata* | plant | 0.88352 | 0.04982 | yes |
| *Daphnopsis schwackeana* | plant | 0.87212 | 0.30708 | yes |
| *Declieuxia deltoidea* | plant | 0.98015 | -0.37617 | no |
| *Delonix regia* | plant | 0.79031 | 0.03354 | yes |
| *Desmodium uncinatum* | plant | 0.77533 | 0.69778 | yes |
| *Diplusodon orbicularis* | plant | 0.98319 | -0.1435 | no |
| *Diplusodon parvifolius* | plant | 0.96570 | -0.37648 | no |
| *Diplusodon ulei* | plant | 0.94503 | -0.23702 | no |

| | | | | |
|---|---|---|---|---|
| *Disynaphia ligulifolia* | plant | 0.87007 | 0.28347 | yes |
| *Duranta erecta* | plant | 0.75165 | 0.15810 | yes |
| *Eichhornia crassipes* | plant | 0.77802 | -0.59863 | no |
| *Epistephium sclerophyllum* | plant | 0.78046 | -0.17661 | no |
| *Eremanthus capitatus* | plant | 0.83456 | -0.03478 | no |
| *Eriope salviifolia* | plant | 0.89543 | -0.10896 | no |
| *Erythrina crista-galli* | plant | 0.72838 | 0.10149 | yes |
| *Eryngium elegans* | plant | 0.81384 | 0.23303 | yes |
| *Eryngium eriophorum* | plant | 0.94075 | 0.00383 | yes |
| *Eryngium horridum* | plant | 0.72755 | 0.33207 | yes |
| *Eryngium sanguisorba* | plant | 0.74961 | 0.22670 | yes |
| *Euphorbia milii* | plant | 0.85601 | -0.65750 | no |
| *Evolvulus glomeratus* | plant | 0.80006 | -0.27981 | no |
| *Faramea montevidensis* | plant | 0.77199 | 0.37995 | yes |
| *Ficus cestrifolia* | plant | 0.75916 | 0.54994 | yes, but isolated in forest-spp only networks |
| *Ficus crocata* | plant | 0.72044 | -0.35764 | no |
| *Ficus obtusiuscula* | plant | 0.73198 | -0.47245 | no |
| *Forsteronia glabrescens* | plant | 0.77180 | -0.05663 | no |
| *Fridericia dispar* | plant | 0.86894 | -0.40768 | no |
| *Galianthe fastigiata* | plant | 0.96901 | 0.41316 | yes |
| *Galianthe valerianoides* | plant | 0.75494 | 0.55848 | yes |
| *Genipa americana* | plant | 0.75449 | -0.49638 | no |
| *Glechon ciliata* | plant | 0.74493 | -0.09383 | no |

| | | | | |
|---|---|---|---|---|
| *Glechon marifolia* | plant | 0.95904 | 0.29288 | yes |
| *Gliricidia sepium* | plant | 0.80613 | -0.22467 | no |
| *Gomphrena demissa* | plant | 0.70322 | -0.25232 | no |
| *Grazielia intermedia* | plant | 0.81772 | 0.48800 | yes |
| *Guatteria australis* | plant | 0.70397 | 0.59693 | yes |
| *Guazuma ulmifolia* | plant | 0.76086 | -0.51866 | no |
| *Habenaria macronectar* | plant | 0.91135 | 0.38111 | yes |
| *Habenaria montevidensis* | plant | 0.98486 | 0.37532 | yes |
| *Handroanthus albus* | plant | 0.77123 | 0.21691 | yes |
| *Heimia salicifolia* | plant | 0.84850 | 0.28494 | yes |
| *Heliconia psittacorum* | plant | 0.73252 | -0.15593 | no |
| *Hennecartia omphalandra* | plant | 0.83989 | 0.35946 | yes |
| *Hillia parasitica* | plant | 0.80760 | 0.27568 | yes |
| *Hippeastrum morelianum* | plant | 0.94559 | -0.13049 | no |
| *Hippeastrum psittacinum* | plant | 0.75704 | 0.44892 | yes |
| *Hippeastrum stylosum* | plant | 0.80191 | -0.22747 | no |
| *Hippocratea volubilis* | plant | 0.78725 | -0.25804 | no |
| *Houlletia brocklehurstiana* | plant | 0.84885 | 0.42667 | yes |
| *Hovenia dulcis* | plant | 0.74754 | 0.35383 | yes |
| *Hymenaea courbaril* | plant | 0.78544 | -0.33815 | no |
| *Hypericum brasiliense* | plant | 0.81112 | 0.34090 | yes |
| *Hyptis lacustris* | plant | 0.92386 | 0.12390 | yes |
| *Hyptis lappulacea* | plant | 0.72358 | 0.61226 | yes |

| | | | | |
|---|---|---|---|---|
| *Hypolytrum schraderianum* | plant | 0.83937 | 0.36921 | yes |
| *Ilex pseudobuxus* | plant | 0.70022 | 0.37880 | yes |
| *Impatiens walleriana* | plant | 0.75261 | 0.42947 | yes |
| *Inga marginata* | plant | 0.77074 | 0.16748 | yes |
| *Inga sessilis* | plant | 0.72047 | 0.72066 | yes |
| *Ipomoea asarifolia* | plant | 0.87329 | -0.45407 | no |
| *Ipomoea bahiensis* | plant | 0.77495 | -0.33173 | no |
| *Ipomoea carnea* | plant | 0.77973 | -0.44497 | no |
| *Ipomoea grandifolia* | plant | 0.82414 | 0.01005 | yes |
| *Ipomoea pes-caprae* | plant | 0.84233 | 0.11131 | yes |
| *Ipomoea purpurea* | plant | 0.76642 | 0.34259 | yes |
| *Jacquemontia nodiflora* | plant | 0.85979 | -0.23576 | no |
| *Jacaranda puberula* | plant | 0.72185 | 0.50530 | yes |
| *Kielmeyera albopunctata* | plant | 0.73103 | 0.14206 | yes |
| *Lagerstroemia indica* | plant | 0.82810 | 0.04253 | yes |
| *Laportea aestuans* | plant | 0.74748 | -0.21370 | no |
| *Leandra australis* | plant | 0.76018 | 0.58383 | yes |
| *Leandra hirtella* | plant | 0.86923 | 0.62599 | yes |
| *Leandra lacunosa* | plant | 0.79771 | 0.21617 | yes |
| *Leandra reversa* | plant | 0.87191 | 0.13254 | yes |
| *Leonurus japonicus* | plant | 0.70205 | 0.26975 | yes |
| *Lessingianthus hypochaeris* | plant | 0.84301 | 0.01745 | yes |
| *Licania dealbata* | plant | 0.96144 | -0.15493 | no |
| *Ludwigia multinervia* | plant | 0.89030 | 0.18090 | yes |
| *Ludwigia peruviana* | plant | 0.75096 | 0.54101 | yes |

| | | | | |
|---|---|---|---|---|
| *Luxemburgia schwackeana* | plant | 0.98156 | -0.18641 | no |
| *Lychnophora salicifolia* | plant | 0.92624 | -0.36479 | no |
| *Macroptilium prostratum* | plant | 0.83799 | 0.07366 | yes |
| *Mandevilla funiformis* | plant | 0.77495 | 0.23302 | yes |
| *Mandevilla hirsuta* | plant | 0.72147 | -0.19249 | no |
| *Mangifera indica* | plant | 0.73861 | -0.50472 | no |
| *Marcgravia polyantha* | plant | 0.83100 | 0.56063 | yes |
| *Miconia cabucu* | plant | 0.85727 | 0.55135 | yes |
| *Miconia cinerascens* | plant | 0.78378 | 0.56925 | yes |
| *Miconia cinnamomifolia* | plant | 0.70851 | 0.40973 | yes |
| *Mikania involucrata* | plant | 0.87482 | 0.44926 | yes |
| *Mikania lundiana* | plant | 0.70288 | 0.27612 | yes |
| *Mikania neurocaula* | plant | 0.98665 | -0.15085 | no |
| *Mikania trinervis* | plant | 0.71022 | 0.53326 | yes |
| *Mimosa arenosa* | plant | 0.82171 | -0.22327 | no |
| *Mimosa aurivillus* | plant | 0.93179 | -0.10575 | no |
| *Mimosa caesalpiniifolia* | plant | 0.82933 | -0.29385 | no |
| *Mimosa pudica* | plant | 0.71381 | -0.28007 | no |
| *Mimosa tenuiflora* | plant | 0.79328 | -0.34454 | no |
| *Mollinedia elegans* | plant | 0.78459 | 0.46246 | yes |
| *Momordica charantia* | plant | 0.74928 | -0.61809 | no |
| *Moquiniastrum cordatum* | plant | 0.97206 | 0.06916 | yes |
| *Muntingia calabura* | plant | 0.79260 | 0.01213 | yes |

| | | | | |
|---|---|---|---|---|
| *Myracrodruon urundeuva* | plant | 0.87251 | -0.39878 | no |
| *Neea schwackeana* | plant | 0.80929 | 0.41677 | yes |
| *Neomitranthes cordifolia* | plant | 0.87922 | 0.02599 | yes |
| *Norantea guianensis* | plant | 0.76857 | -0.25275 | no |
| *Nymphoides indica* | plant | 0.75813 | -0.53326 | no |
| *Ocimum gratissimum* | plant | 0.73219 | -0.07099 | no |
| *Ocotea lancifolia* | plant | 0.72373 | -0.03595 | no |
| *Opuntia monacantha* | plant | 0.73537 | -0.13840 | no |
| *Oreopanax capitatus* | plant | 0.80214 | 0.36432 | yes |
| *Ossaea amygdaloides* | plant | 0.73113 | 0.59596 | yes |
| *Ouratea cuspidata* | plant | 0.71377 | -0.19220 | no |
| *Ouratea hexasperma* | plant | 0.75940 | -0.04072 | no |
| *Ouratea parviflora* | plant | 0.81108 | 0.64960 | yes |
| *Palicourea guianensis* | plant | 0.74073 | 0.28554 | yes |
| *Palicourea marcgravii* | plant | 0.71352 | -0.03081 | no |
| *Palicourea rigida* | plant | 0.85067 | -0.04868 | no |
| *Paliavana tenuiflora* | plant | 0.74526 | 0.01688 | yes |
| *Parthenium hysterophorus* | plant | 0.84824 | -0.47339 | no |
| *Passiflora cincinnata* | plant | 0.80348 | -0.40305 | no |
| *Passiflora morifolia* | plant | 0.70672 | 0.20886 | yes |
| *Paullinia elegans* | plant | 0.72318 | -0.50173 | no |
| *Pavonia cancellata* | plant | 0.74984 | -0.37083 | no |
| *Pavonia friesii* | plant | 0.81306 | 0.25820 | yes |

| | | | | |
|---|---|---|---|---|
| *Pavonia sepium* | plant | 0.74938 | 0.35720 | yes |
| *Paypayrola blanchetiana* | plant | 0.84494 | 0.03483 | yes |
| *Pelexia oestrifera* | plant | 0.77876 | -0.18851 | no |
| *Pfaffia tuberosa* | plant | 0.86464 | 0.19305 | yes |
| *Piper arboreum* | plant | 0.72459 | 0.09433 | yes |
| *Piptocarpha quadrangularis* | plant | 0.80375 | 0.13237 | yes |
| *Piptadenia stipulacea* | plant | 0.81656 | -0.19697 | no |
| *Piptadenia viridiflora* | plant | 0.90678 | -0.30809 | no |
| *Pleroma mutabile* | plant | 0.79475 | 0.25808 | yes |
| *Pleroma trichopodum* | plant | 0.78101 | 0.38013 | yes |
| *Pogonophora schomburgkiana* | plant | 0.72255 | -0.07458 | no |
| *Portulaca oleracea* | plant | 0.78110 | -0.77252 | no |
| *Protium aracouchini* | plant | 0.70675 | 0.09430 | yes |
| *Prosopis juliflora* | plant | 0.93220 | -0.31601 | no |
| *Pseudobrickellia brasiliensis* | plant | 0.92492 | -0.27840 | no |
| *Pseudostifftia kingii* | plant | 0.95027 | -0.58521 | no |
| *Psychotria brachyceras* | plant | 0.90508 | 0.19776 | yes |
| *Psychotria schlechtendaliana* | plant | 0.93178 | 0.22325 | yes |
| *Psychotria suterella* | plant | 0.75166 | 0.64064 | yes |
| *Randia armata* | plant | 0.69996 | -0.54301 | no |
| *Raphanus sativus* | plant | 0.77512 | -0.06934 | no |
| *Rauvolfia grandiflora* | plant | 0.70837 | 0.14063 | yes |
| *Remijia ferruginea* | plant | 0.92444 | -0.07809 | no |
| *Rhododendron* | plant | 0.78595 | 0.09579 | yes |

| | | | | |
|---|---|---|---|---|
| *simsii* | | | | |
| *Rhynchosia corylifolia* | plant | 0.90333 | 0.29269 | yes |
| *Ricinus communis* | plant | 0.72093 | -0.01688 | no |
| *Richardia grandiflora* | plant | 0.72305 | -0.38636 | no |
| *Rubus rosifolius* | plant | 0.76686 | 0.72749 | yes |
| *Rudgea jasminoides* | plant | 0.75384 | 0.49408 | yes |
| *Ruellia affinis* | plant | 0.80660 | 0.18314 | yes |
| *Sabicea villosa* | plant | 0.77405 | 0.40697 | yes |
| *Sagittaria montevidensis* | plant | 0.74940 | 0.26864 | yes |
| *Sarcoglottis fasciculata* | plant | 0.71931 | 0.13636 | yes |
| *Schlechtendalia luzulifolia* | plant | 0.75530 | 0.28521 | yes |
| *Schlumbergera truncata* | plant | 0.83379 | 0.34806 | yes |
| *Sclerolobium aureum* | plant | 0.95294 | -0.25673 | no |
| *Senecio brasiliensis* | plant | 0.76658 | 0.45565 | yes |
| *Senna cana* | plant | 0.90949 | -0.1794 | no |
| *Senna corymbosa* | plant | 0.84953 | 0.02899 | yes |
| *Senecio crassiflorus* | plant | 0.92120 | -0.13625 | no |
| *Senna georgica* | plant | 0.75926 | -0.10466 | no |
| *Senna uniflora* | plant | 0.89750 | -0.20557 | no |
| *Serjania lethalis* | plant | 0.77465 | -0.14978 | no |
| *Sisyrinchium palmifolium* | plant | 0.80491 | 0.36149 | yes |
| *Sisyrinchium vaginatum* | plant | 0.77949 | 0.21760 | yes |
| *Solidago chilensis* | plant | 0.75309 | 0.39524 | yes |
| *Solanum concinnum* | plant | 0.71720 | 0.09079 | yes |

| | | | | |
|---|---|---|---|---|
| *Solanum guaraniticum* | plant | 0.76449 | 0.45007 | yes |
| *Solanum mauritianum* | plant | 0.70509 | 0.30610 | yes |
| *Solanum paludosum* | plant | 0.85042 | -0.30721 | no |
| *Solanum piluliferum* | plant | 0.74911 | 0.64464 | yes |
| *Solanum rufescens* | plant | 0.70306 | 0.36412 | yes |
| *Spondias mombin* | plant | 0.87604 | -0.13966 | no |
| *Stachytarpheta maximiliani* | plant | 0.77314 | 0.03743 | yes |
| *Stenocephalum megapotamicum* | plant | 0.81929 | 0.16836 | yes |
| *Stigmaphyllon paralias* | plant | 0.75844 | -0.16850 | no |
| *Stryphnodendron pulcherrimum* | plant | 0.81941 | -0.15305 | no |
| *Styrax camporum* | plant | 0.74435 | -0.19626 | no |
| *Stylosanthes montevidensis* | plant | 0.75929 | -0.21141 | no |
| *Syagrus coronata* | plant | 0.77873 | -0.39047 | no |
| *Symphyopappus reticulatus* | plant | 0.87367 | -0.13030 | no |
| *Syzygium cumini* | plant | 0.76619 | -0.44050 | no |
| *Tecoma ype* | plant | 0.93806 | -0.12721 | no |
| *Temnadenia odorifera* | plant | 0.78171 | -0.21800 | no |
| *Tetrapterys microphylla* | plant | 0.96450 | -0.10326 | no |
| *Tibouchina barnebyana* | plant | 0.93381 | -0.01248 | no |
| *Tibouchina cerastifolia* | plant | 0.85200 | 0.62601 | yes |
| *Tibouchina clavata* | plant | 0.72896 | 0.30705 | yes |
| *Tibouchina regnellii* | plant | 0.87477 | 0.27917 | yes |
| *Tibouchina* | plant | 0.90430 | 0.55090 | yes |

| | | | | |
|---|---|---|---|---|
| *urvilleana* | | | | |
| *Tipuana tipu* | plant | 0.86403 | 0.38534 | yes |
| *Tocoyena bullata* | plant | 0.80314 | -0.28002 | no |
| *Tripodanthus acutifolius* | plant | 0.71268 | 0.14757 | yes |
| *Trixis praestans* | plant | 0.79125 | 0.37117 | yes |
| *Tropaeolum majus* | plant | 0.83172 | 0.20185 | yes |
| *Turnera subulata* | plant | 0.83957 | -0.42338 | no |
| *Vernonanthura beyrichii* | plant | 0.74099 | 0.25294 | yes |
| *Verbena bonariensis* | plant | 0.77399 | 0.39670 | yes |
| *Vernonanthura discolor* | plant | 0.76797 | 0.72433 | yes |
| *Verbena litoralis* | plant | 0.71221 | 0.41464 | yes |
| *Vernonanthura nudiflora* | plant | 0.83294 | 0.13043 | yes |
| *Vernonanthura puberula* | plant | 0.84033 | 0.64454 | yes |
| *Vernonanthura tweediana* | plant | 0.73834 | 0.34263 | yes |
| *Vitex triflora* | plant | 0.90901 | -0.12143 | no |
| *Vriesea gigantea* | plant | 0.80960 | 0.47646 | yes |
| *Vriesea hoehneana* | plant | 0.88188 | 0.18567 | yes |
| *Vriesea philippocoburgii* | plant | 0.79765 | 0.41983 | yes |
| *Vriesea vagans* | plant | 0.83194 | 0.61815 | yes |
| *Waltheria indica* | plant | 0.71933 | -0.41773 | no |
| *Agraulis vanillae* | insect pollinator | 0.72259 | -0.27653 | no |
| *Anidarnes dissidens* | insect pollinator | 0.81163 | -0.32422 | no |
| *Ascia monuste* | insect pollinator | 0.80314 | -0.30828 | no |
| *Atta sexdens* | insect pollinator | 0.84054 | -0.08607 | no |
| *Augochloropsis brachycephala* | insect pollinator | 0.99271 | 0.07943 | yes |

| | | | | |
|---|---|---|---|---|
| *Augochlora cephalica* | insect pollinator | 0.99220 | 0.06553 | yes |
| *Augochlorella ephyra* | insect pollinator | 0.97667 | 0.46926 | yes |
| *Augochloropsis patens* | insect pollinator | 0.99271 | 0.07932 | yes |
| *Battus polydamas* | insect pollinator | 0.93187 | 0.19068 | yes |
| *Bombus brasiliensis* | insect pollinator | 0.84491 | 0.47514 | yes |
| *Bombus brevivillus* | insect pollinator | 0.78455 | -0.08989 | no |
| *Bombus pauloensis* | insect pollinator | 0.91603 | 0.30396 | yes |
| *Brachygastra lecheguana* | insect pollinator | 0.71290 | 0.34950 | yes |
| *Callimormus corades* | insect pollinator | 0.99297 | 0.08428 | yes |
| *Camponotus crassus* | insect pollinator | 0.93112 | 0.05447 | yes |
| *Camponotus rufipes* | insect pollinator | 0.78610 | 0.47056 | yes |
| *Centris analis* | insect pollinator | 0.75910 | -0.15344 | no |
| *Centris flavifrons* | insect pollinator | 0.71805 | -0.29831 | no |
| *Centris fuscata* | insect pollinator | 0.85188 | -0.45745 | no |
| *Centris varia* | insect pollinator | 0.77988 | -0.66903 | no |
| *Cephalotes atratus* | insect pollinator | 0.77644 | -0.17055 | no |
| *Cephalotes pusillus* | insect pollinator | 0.87280 | -0.17041 | no |
| *Cycloneda sanguinea* | insect pollinator | 0.90838 | 0.26059 | yes |
| *Diabrotica speciosa* | insect pollinator | 0.82469 | -0.06836 | no |
| *Drosophila bromelioides* | insect pollinator | 0.81516 | 0.44855 | yes |
| *Epicharis analis* | insect pollinator | 0.70576 | 0.02852 | yes |
| *Euglossa annectans* | insect pollinator | 0.85906 | 0.14865 | yes |
| *Euglossa cordata* | insect pollinator | 0.85543 | 0.27680 | yes |
| *Euglossa fimbriata* | insect pollinator | 0.71153 | -0.18494 | no |
| *Euglossa pleosticta* | insect pollinator | 0.75032 | 0.08047 | yes |

| | | | | |
|---|---|---|---|---|
| *Eulaema nigrita* | insect pollinator | 0.76774 | -0.2979 | no |
| *Exaerete smaragdina* | insect pollinator | 0.82447 | 0.09985 | yes |
| *Exomalopsis analis* | insect pollinator | 0.99566 | 0.00349 | yes |
| *Exomalopsis auropilosa* | insect pollinator | 0.96760 | -0.27562 | no |
| *Geotrigona subterranea* | insect pollinator | 0.92428 | -0.06214 | no |
| *Gnamptogenys striatula* | insect pollinator | 0.79941 | 0.23239 | yes |
| *Heliopetes arsalte* | insect pollinator | 0.79685 | 0.00809 | yes |
| *Heteroponera dolo* | insect pollinator | 0.72794 | 0.12526 | yes |
| *Hylomyrma balzani* | insect pollinator | 0.92184 | -0.09449 | no |
| *Hypoponera foreli* | insect pollinator | 0.91310 | -0.05346 | no |
| *Idarnes brevicollis* | insect pollinator | 0.71433 | -0.21510 | no |
| *Idarnes carme* | insect pollinator | 0.74934 | -0.34640 | no |
| *Idarnes dimorphicus* | insect pollinator | 0.87475 | -0.26865 | no |
| *Idarnes maximus* | insect pollinator | 0.73941 | -0.25065 | no |
| *Idarnes punctatus* | insect pollinator | 0.84106 | -0.25438 | no |
| *Lachnomyrmex plaumanni* | insect pollinator | 0.94409 | 0.78858 | yes |
| *Lachnomyrmex victori* | insect pollinator | 0.86412 | 0.56433 | yes |
| *Linepithema iniquum* | insect pollinator | 0.98928 | -0.01735 | no |
| *Linepithema neotropicum* | insect pollinator | 0.81834 | -0.12455 | no |
| *Mechanitis lysimnia* | insect pollinator | 0.72276 | 0.14657 | yes |
| *Megachile susurrans* | insect pollinator | 0.81808 | 0.11803 | yes |
| *Melipona mondury* | insect pollinator | 0.81798 | 0.09714 | yes |
| *Melipona quadrifasciata* | insect pollinator | 0.71797 | 0.22771 | yes |
| *Melipona* | insect pollinator | 0.86867 | 0.30997 | yes |

| | | | | |
|---|---|---|---|---|
| *rufiventris* | | | | |
| *Melitoma segmentaria* | insect pollinator | 0.92058 | -0.18392 | no |
| *Mischocyttarus drewseni* | insect pollinator | 0.99559 | 0.21734 | yes |
| *Mischocyttarus rotundicollis* | insect pollinator | 0.92058 | 0.12047 | yes |
| *Myrmelachista catharinae* | insect pollinator | 0.78538 | -0.05236 | no |
| *Oxaea flavescens* | insect pollinator | 0.90538 | -0.37693 | no |
| *Palpada conica* | insect pollinator | 0.98849 | -0.08938 | no |
| *Paratetrapedia fervida* | insect pollinator | 0.82119 | 0.08017 | yes |
| *Partamona helleri* | insect pollinator | 0.73772 | 0.12518 | yes |
| *Paratrigona subnuda* | insect pollinator | 0.77169 | 0.42119 | yes |
| *Pegoscapus aerumnosus* | insect pollinator | 0.91761 | -0.19060 | no |
| *Phoebis argante* | insect pollinator | 0.78953 | 0.36623 | yes |
| *Plebeia emerina* | insect pollinator | 0.95423 | 0.21821 | yes |
| *Plebeia remota* | insect pollinator | 0.74103 | 0.52236 | yes |
| *Polybia ignobilis* | insect pollinator | 0.73384 | 0.26479 | yes |
| *Polybia sericea* | insect pollinator | 0.82129 | -0.09453 | no |
| *Protambulyx strigilis* | insect pollinator | 0.70645 | 0.36470 | yes |
| *Pseudodorus clavatus* | insect pollinator | 0.82096 | 0.32442 | yes |
| *Ptilothrix tricolor* | insect pollinator | 0.99471 | -0.00701 | no |
| *Salpingogaster nigra* | insect pollinator | 0.99128 | 0.05880 | yes |
| *Scaptotrigona xanthotricha* | insect pollinator | 0.99127 | 0.05424 | yes |
| *Schwarziana quadripunctata* | insect pollinator | 0.71250 | 0.42710 | yes |
| *Tegosa claudina* | insect pollinator | 0.71911 | 0.16479 | yes |
| *Thygater analis* | insect pollinator | 0.77471 | 0.16495 | yes |

| | | | | |
|---|---|---|---|---|
| *Toxomerus basalis* | insect pollinator | 0.99433 | 0.06937 | yes |
| *Toxomerus norma* | insect pollinator | 0.99201 | 0.08326 | yes |
| *Toxomerus pictus* | insect pollinator | 0.99271 | 0.07883 | yes |
| *Toxomerus productus* | insect pollinator | 0.99307 | 0.08139 | yes |
| *Toxomerus steatogaster* | insect pollinator | 0.99307 | 0.08191 | yes |
| *Toxomerus virgulatus* | insect pollinator | 0.98892 | 0.01093 | yes |
| *Trigona braueri* | insect pollinator | 0.94909 | 0.37908 | yes |
| *Trina geometrina* | insect pollinator | 0.78558 | 0.43088 | yes |
| *Urbanus simplicius* | insect pollinator | 0.79949 | -0.06998 | no |
| *Urbanus teleus* | insect pollinator | 0.82502 | 0.35825 | yes |
| *Vanessa braziliensis* | insect pollinator | 0.78495 | 0.47514 | yes |
| *Vanessa myrinna* | insect pollinator | 0.72604 | -0.00269 | no |
| *Xylocopa brasilianorum* | insect pollinator | 0.95362 | 0.10404 | yes |
| *Xylocopa ciliata* | insect pollinator | 0.99663 | 0.17876 | yes |
| *Xylocopa suspecta* | insect pollinator | 0.74791 | 0.27223 | yes |

**Table 3.**

Minimum and maximum values of the studied network properties for all-spp and forest-spp networks, with sites of 25km N-S length.

| | **all-spp** | | **forest-spp** | |
|---|---|---|---|---|
| **network properties** | **min** | **max** | **min** | **max** |
| $N_{pl}$ | 273 | 297 | 159 | 163 |
| $N_{po}$ | 66 | 92 | 42 | 58 |
| L | 3995 | 5648 | 1803 | 2587 |
| SLW | 1.33211 | 42.17379 | 0.22008 | 38.57559 |
| $HS_{pl}$ | 2.40993 | 4.1966 | 1.55808 | 3.74466 |
| $HS_{po}$ | 1.47311 | 3.69256 | 1.06112 | 3.40528 |
| C | 0.00712 | 0.02115 | 0.00814 | 0.03464 |
| Centr | 0.75128 | 0.79556 | 0.66419 | 0.73824 |
| A | -0.11582 | 0.03233 | -0.08182 | 0.01404 |
| M | 0.31453 | 0.80177 | 0.31584 | 0.7904 |
| $Nb_{mod}$ | 6 | 29 | 5 | 22 |
| Nest | 0.09774 | 2.90643 | 0.02969 | 2.77831 |

### Table 4.

Spearman's rho correlation index between network properties and forest or land cover heterogeneity (HLC) for all-spp and forest-spp networks. The symbols show positive i.e. $\rho > 0.3$ (▲), negative i.e. $\rho < -0.3$ (▼), weak positive i.e. $0.1 < \rho < 0.3$ (■▴), weak negative i.e. $-0.3 < \rho < -0.1$ (■▾) or no correlation i.e. $-0.1 < \rho < 0.1$ or p-value > 0.05 (■). Rho values are the average of 10 repetitions using the `jitter` function in R for both variables.

| all-spp | | $N_{pl}$ | $N_{po}$ | L | SLW | $HS_{pl}$ | $HS_{po}$ | C | Centr | M | $Nb_{mod}$ | A | Nest |
|---|---|---|---|---|---|---|---|---|---|---|---|---|---|
| forest cover | 25 km | -0.32 ▼ | -0.37 ▼ | -0.38 ▼ | 0.2 ■▴ | 0.12 ■▴ | 0.32 ▲ | 0.26 ■▴ | 0.36 ▲ | -0.15 ■▾ | n.s. ■ | -0.06 ■ | 0.14 ■▴ |
| | 50 km | -0.36 ▼ | -0.37 ▼ | -0.38 ▼ | 0.26 ■▴ | 0.09 ■ | 0.31 ▲ | 0.19 ■▴ | 0.37 ▲ | -0.12 ■▾ | n.s. ■ | n.s. ■ | 0.21 ■▴ |
| | 100 km | -0.36 ▼ | -0.31 ▼ | -0.33 ▼ | 0.36 ▲ | n.s. ■ | 0.33 ▲ | n.s. ■ | 0.34 ■▴ | n.s. ■ | n.s. ■ | n.s. ■ | 0.30 ■▴ |
| HLC | 25 km | -0.08 ■ | -0.42 ▼ | -0.41 ▼ | -0.18 ■▾ | n.s. ■ | 0.12 ■▴ | 0.05 ■ | 0.42 ▲ | n.s. ■ | n.s. ■ | n.s. ■ | -0.17 ■▾ |
| | 50 km | -0.12 ■▾ | -0.45 ▼ | -0.44 ▼ | -0.14 ■▾ | n.s. ■ | n.s. ■ | n.s. ■ | 0.46 ▲ | n.s. ■ | n.s. ■ | n.s. ■ | -0.09 ■ |
| | 100 km | n.s. ■ | -0.52 ▼ | -0.47 ▼ | n.s. ■ | n.s. ■ | n.s. ■ | n.s. ■ | 0.50 ▲ | n.s. ■ | n.s. ■ | n.s. ■ | n.s. ■ |

| forest-spp | | $N_{pl}$ | $N_{po}$ | L | SLW | $HS_{pl}$ | $HS_{po}$ | C | Centr | M | $Nb_{mod}$ | A | Nest |
|---|---|---|---|---|---|---|---|---|---|---|---|---|---|
| forest cover | 25 km | n.s. ■ | -0.27 ■▾ | -0.25 ■▾ | 0.71 ▲ | 0.57 ▲ | 0.47 ▲ | 0.39 ▲ | 0.21 ■▴ | 0.17 ■▴ | n.s. ■ | -0.57 ▼ | 0.59 ▲ |
| | 50 km | n.s. ■ | -0.28 ■▾ | -0.27 ■▾ | 0.78 ▲ | 0.58 ▲ | 0.54 ▲ | 0.34 ▲ | 0.24 ■▴ | 0.18 ■▴ | 0.10 ■ | -0.59 ▼ | 0.66 ▲ |
| | 100 km | n.s. ■ | n.s. ■▾ | n.s. ▼ | 0.85 ▲ | 0.60 ▲ | 0.66 ▲ | 0.27 ■▴ | n.s. ■▴ | n.s. ■ | n.s. ■ | -0.63 ▼ | 0.73 ▲ |

| | | | | | | | | | | | | | |
|---|---|---|---|---|---|---|---|---|---|---|---|---|---|
| HLC | 25 km | n.s. | -0.41 | -0.41 | 0.15 | 0.20 | 0.12 | -0.09 | 0.40 | 0.26 | 0.06 | -0.18 | 0.10 |
| | | ■ | ▼ | ▼ | ■▴ | ■▴ | ■▴ | ■ | ▲ | ■▴ | ■ | ■▾ | ■ |
| | 50 km | n.s. | -0.47 | -0.47 | 0.18 | 0.16 | 0.13 | -0.17 | 0.47 | 0.26 | n.s. | -0.11 | 0.16 |
| | | ■ | ▼ | ▼ | ■▴ | ■▴ | ■▴ | ■▾ | ▲ | ■▴ | ■ | ■▴ | ■▴ |
| | 100 km | n.s. | -0.54 | -0.57 | 0.21 | n.s. | n.s. | -0.26 | 0.54 | 0.24 | n.s. | n.s. | n.s. |
| | | ■ | ▼ | ▼ | ■▴ | ■ | ■ | ■▾ | ▲ | ■▴ | ■ | ■ | ■ |

### Figure 1.

**Blockmodels of networks.** Example of blockmodel for a hypothetical network where white flowers usually are visited by moths, and never by butterflies, pink flowers are visited by both butterflies and bees, and yellow and blue flowers are visited only by bees. The species are partitioned into six groups, and we visualize the block model as a block matrix, where each entry $b_{ij}$ corresponds to the probability of a link between a node of group $i$ and a node of group $j$. Darker colors indicate higher probability.

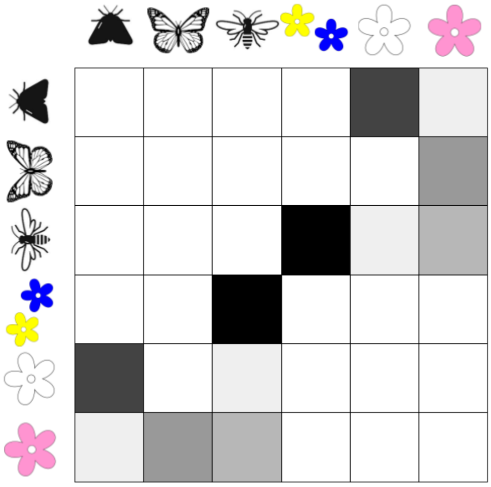

### Figure 2.

**Network properties and habitat variables**. Each plot shows the corresponding Spearman's rank correlation.

All-spp networks: Network properties ~ forest cover (25 km scale)

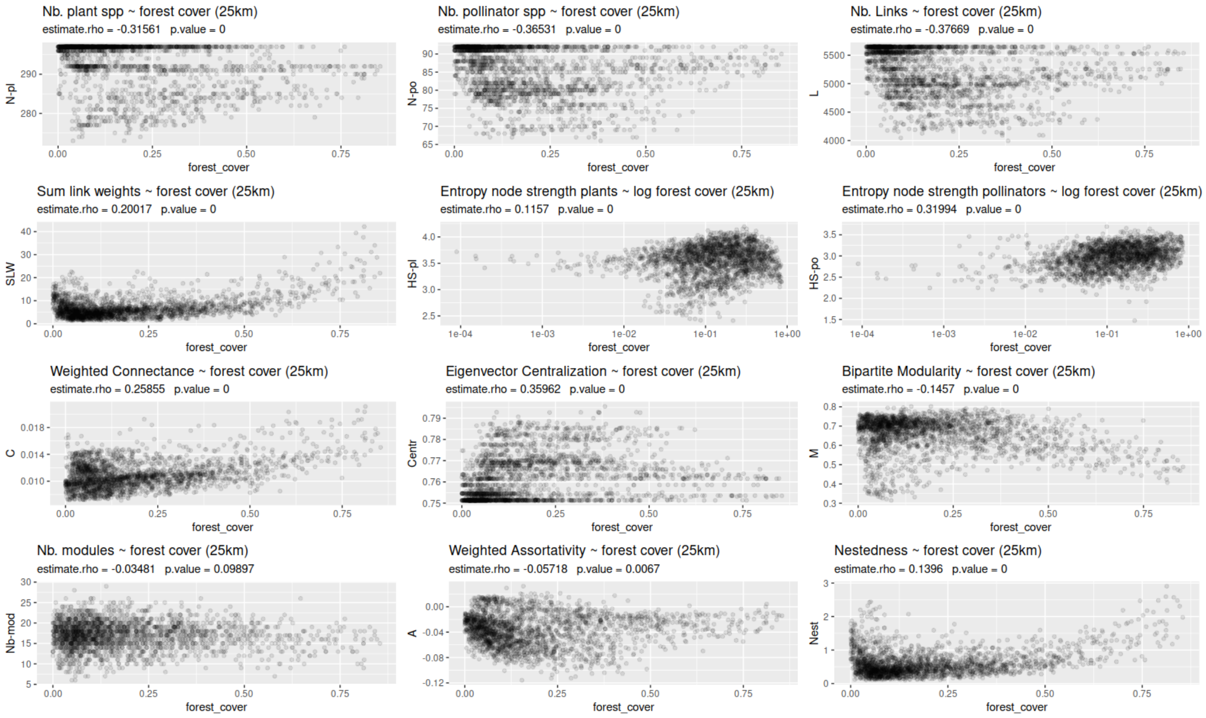


All-spp networks: Network properties ~ land cover heterogeneity (25 km scale)

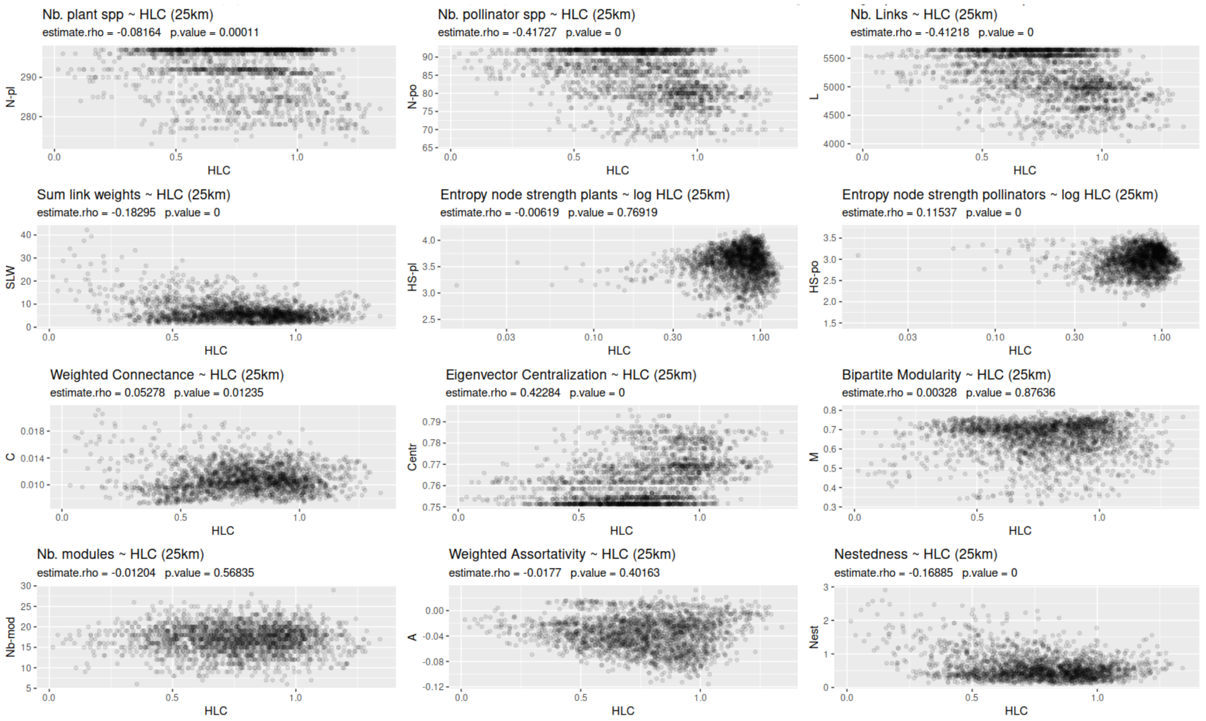

### All-spp networks: Network properties ~ forest cover (50 km scale)

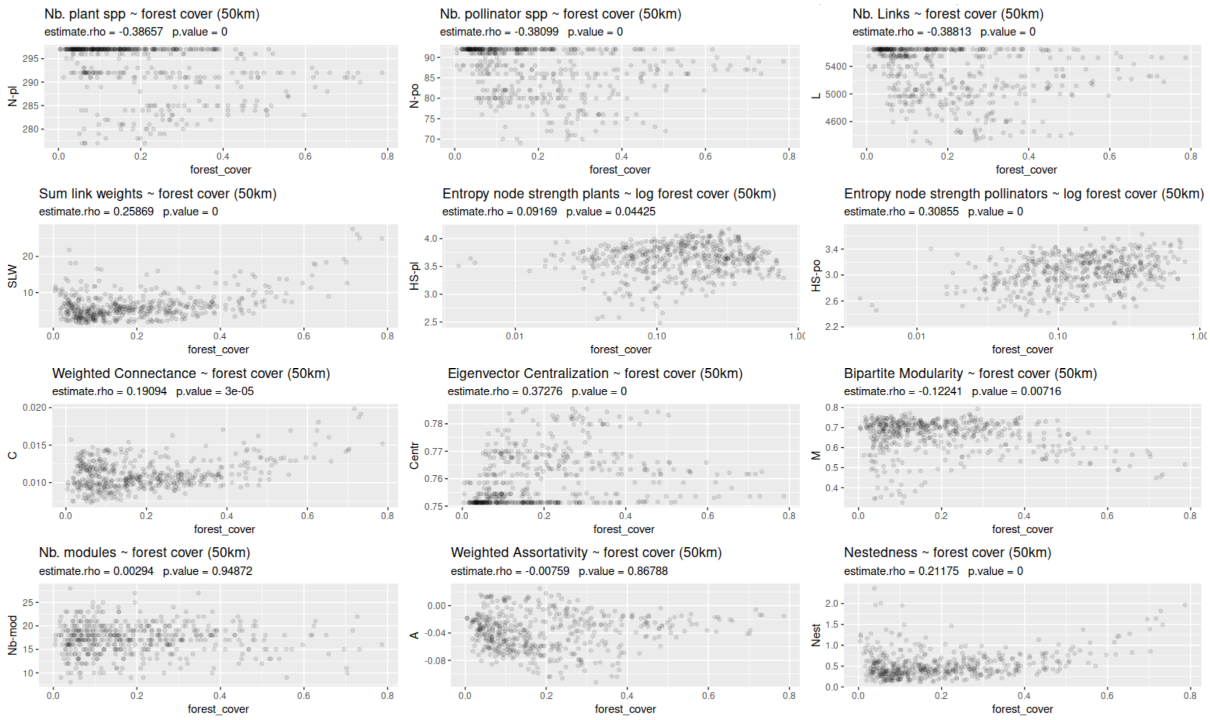


### All-spp networks: Network properties ~ land cover heterogeneity (50 km scale)

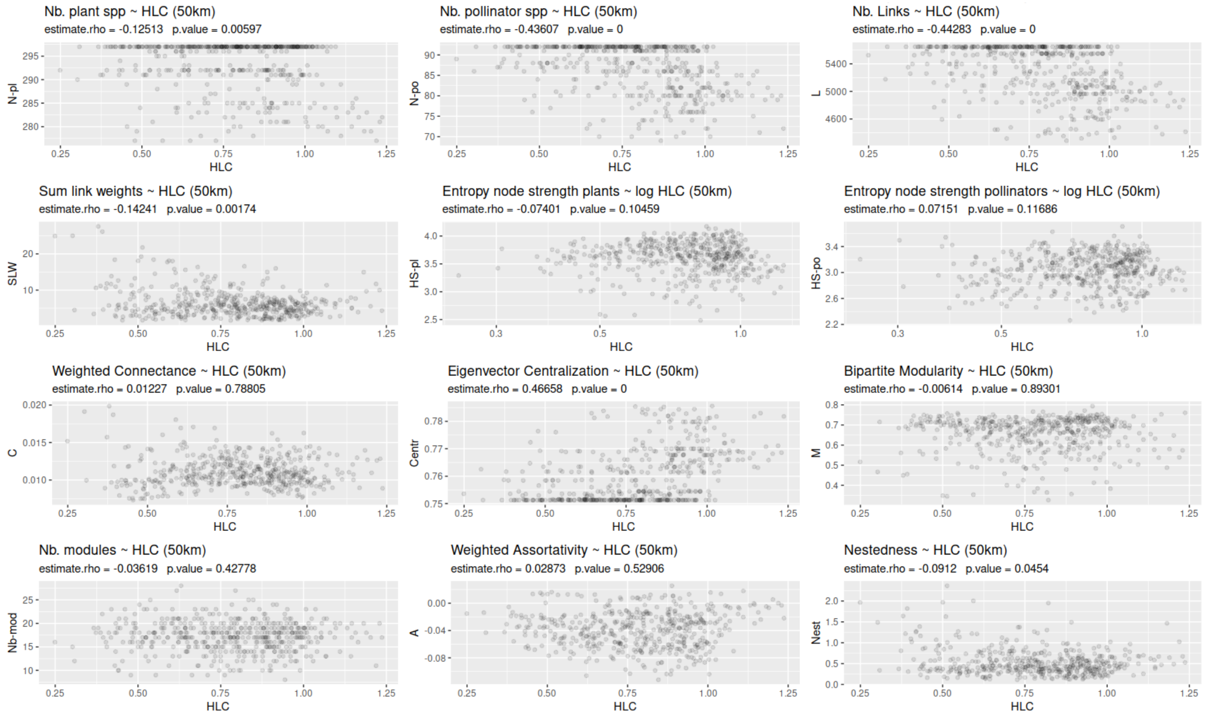

### All-spp networks: Network properties ~ forest cover (100 km scale)

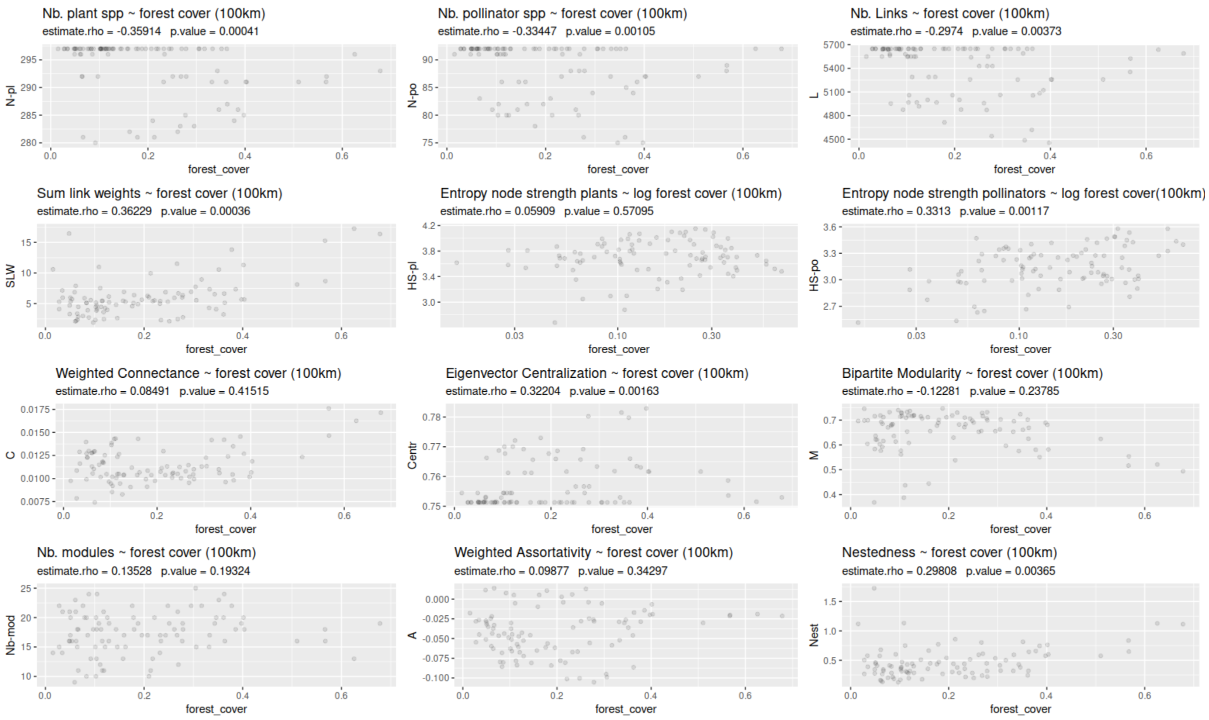


### All-spp networks: Network properties ~ land cover heterogeneity (100 km scale)

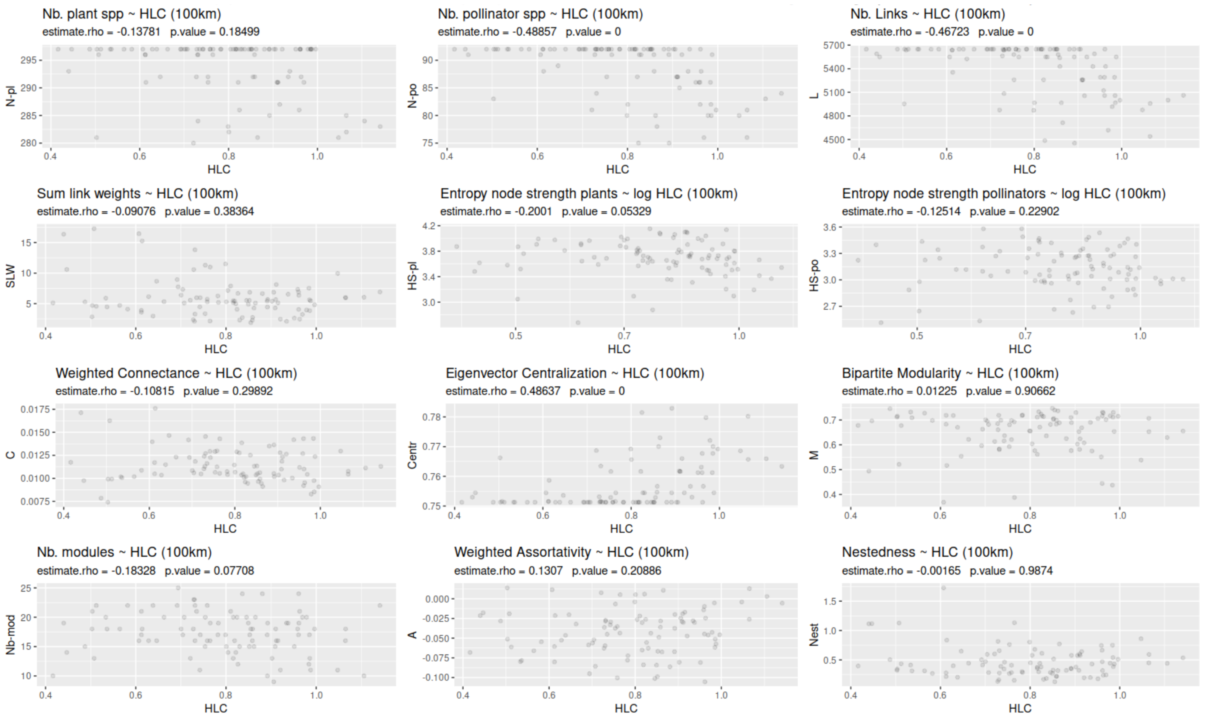

### Forest-spp networks: Network properties ~ forest cover (25 km scale)

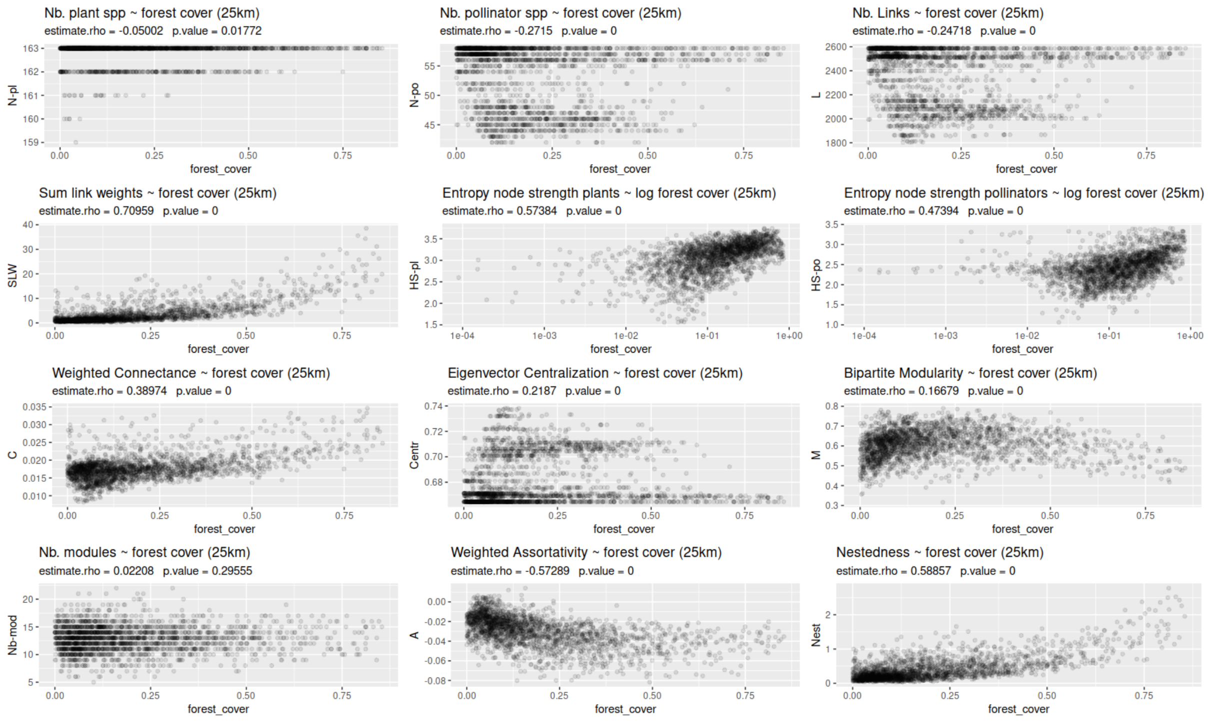


### Forest-spp networks: Network properties ~ land cover heterogeneity (25 km scale)

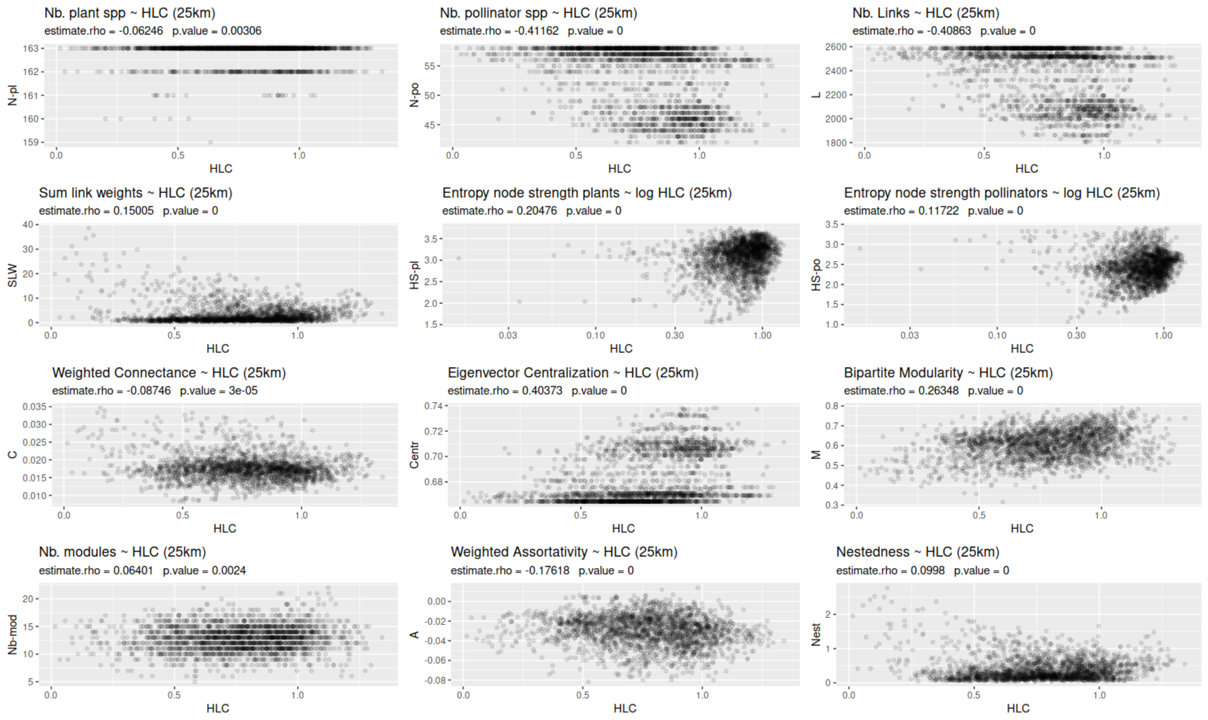

### Forest-spp networks: Network properties ~ forest cover (50 km scale)

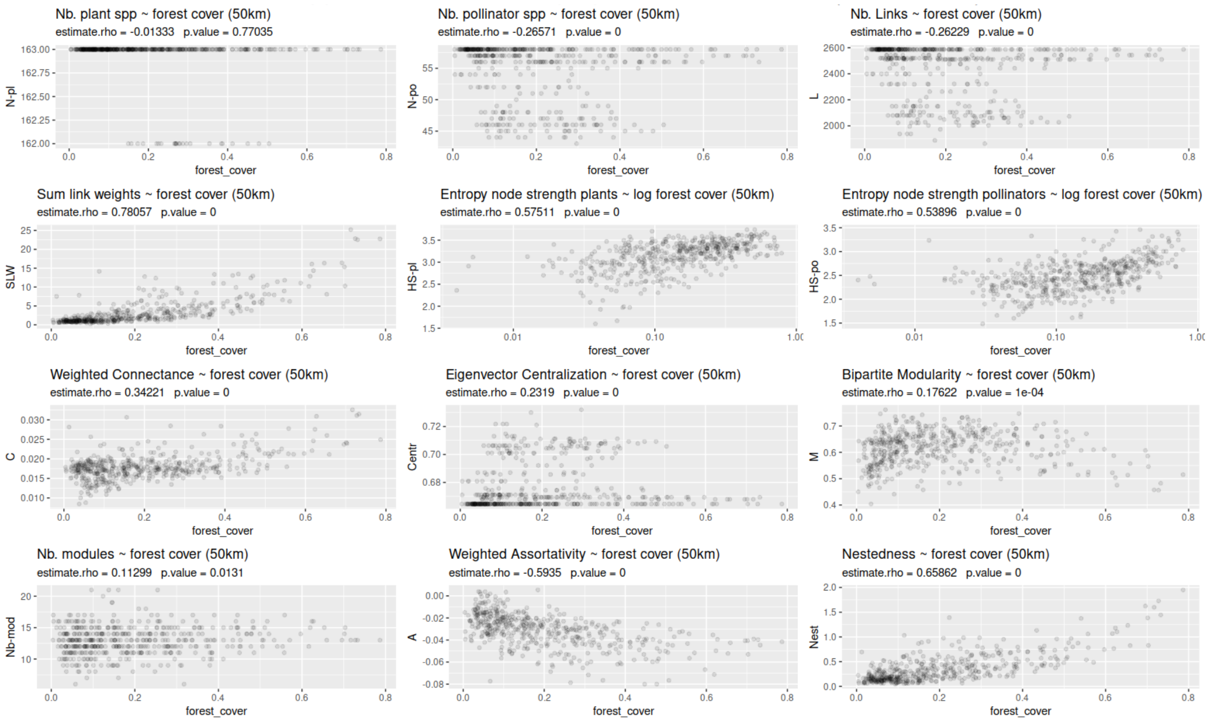


### Forest-spp networks: Network properties ~ land cover heterogeneity (50 km scale)

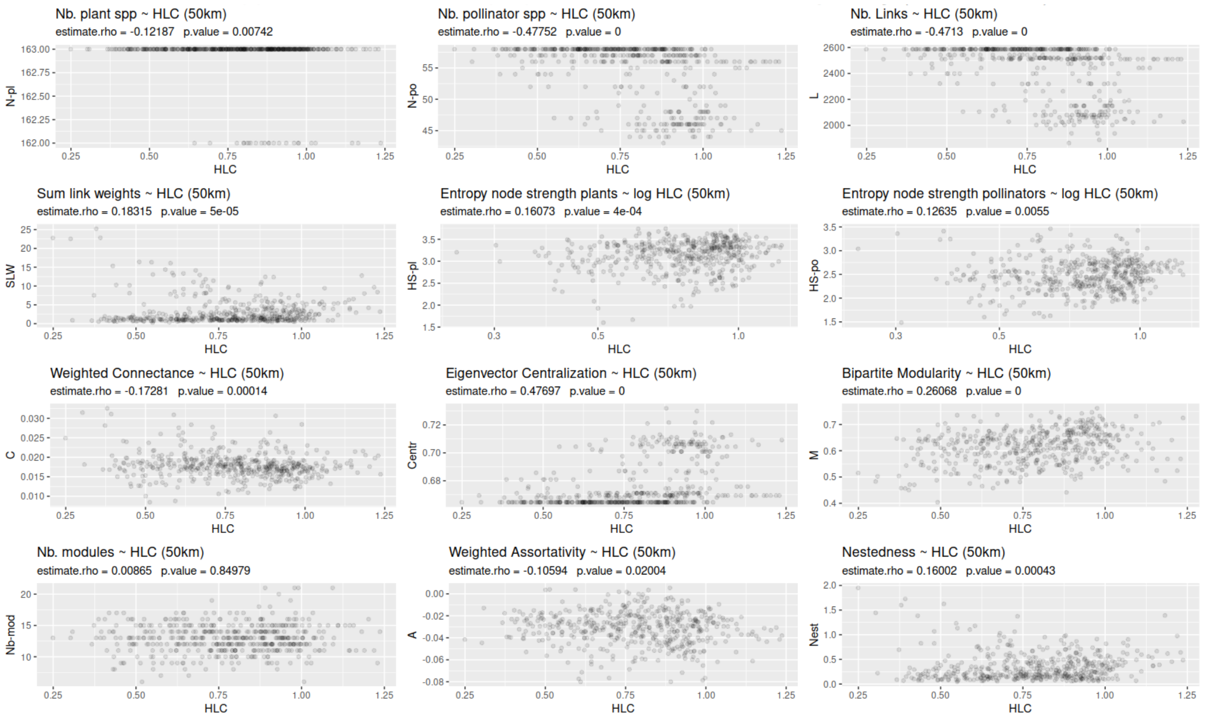

### Forest-spp networks: Network properties ~ forest cover (100 km scale)

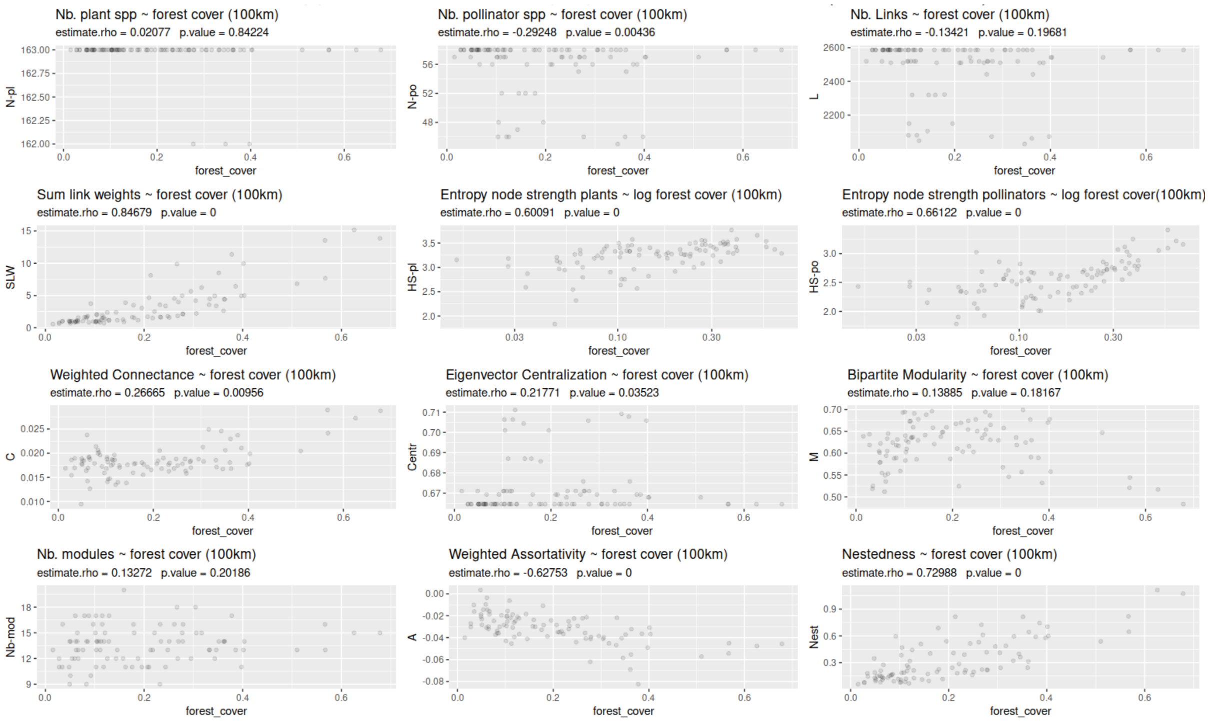


### Forest-spp networks: Network properties ~ land cover heterogeneity (100 km scale)

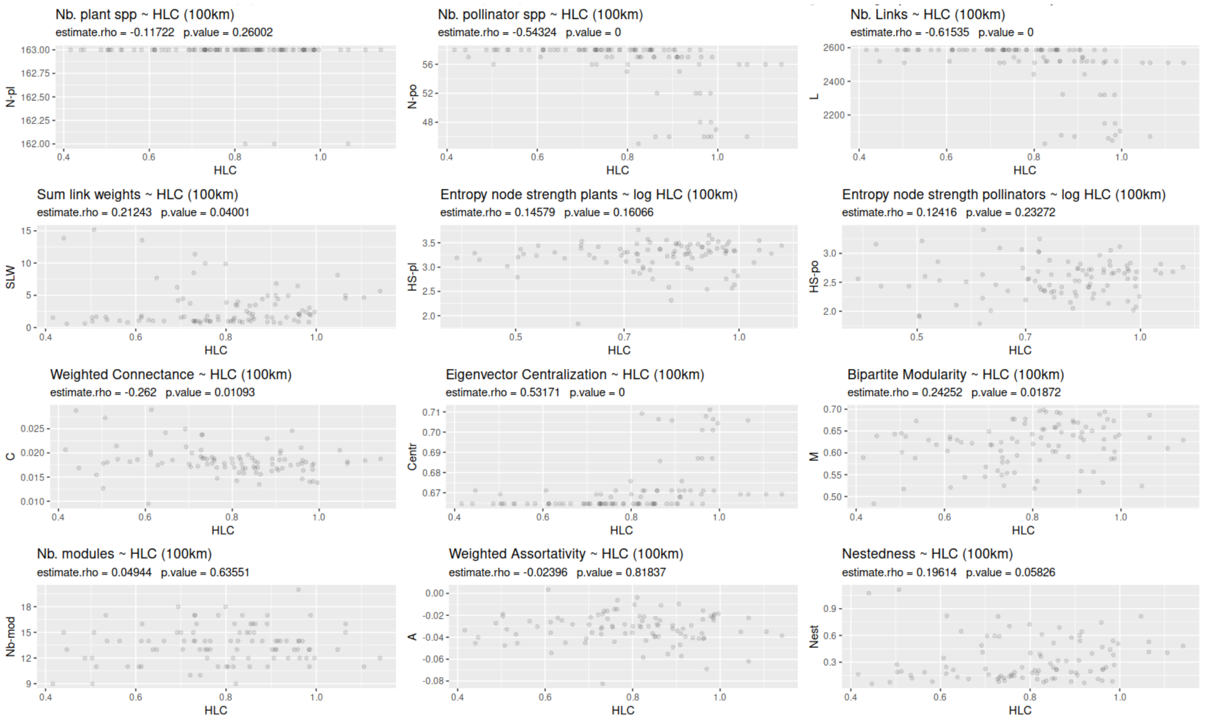

### Figure 3.

Three dimensional plot showing forest cover, number of links (L) and sum of link weights (SLW) for the forest-spp networks in the 25 km scale. Networks of high forest cover sites are always relatively large, both in L and N (not shown) and have higher S, while those sites with little forest show a wide range of network sizes (L), but their total strength S is always low. We see that only high forest sites have networks with higher S, while L is independent of forest cover.

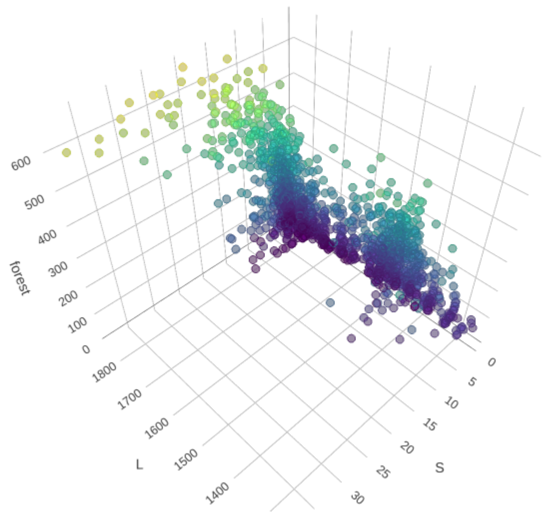
